# Phages communicate across species to shape microbial ecosystems

**DOI:** 10.1101/2025.10.13.681337

**Authors:** Francisca Gallego-del-Sol, Daniel Sin, Cora Chmielowska, Javier Mancheño-Bonillo, Yuyi Li, Sara Zamora-Caballero, Nuria Quiles-Puchalt, José R Penadés, Alberto Marina

**Author notes:** Corresponding authors: Alberto Marina, Instituto de Biomedicina de Valencia, Spain, José R Penadés, Centre for Bacterial Resistance Biology, Imperial College London, UK. These authors contributed equally (Francisca Gallego-del-Sol, Daniel Sin, Cora Chmielowska).

## Abstract

Arbitrium is a communication system that helps bacteriophages decide between lysis and lysogeny via secreted peptides. In arbitrium, the AimP peptide binds its cognate AimR receptor to repress *aimX* expression, promoting lysogeny. It has been assumed that each AimR responds exclusively to its own AimP. Here, we question this view by demonstrating cross-communication between distinct arbitrium systems. Using prototypical arbitrium phages, we demonstrate that AimP peptides bind and repress unrelated AimR receptors, promoting lysogeny and reducing prophage induction. Structural and binding assays reveal conserved residues enabling cross-recognition while preserving specificity. In mixed lysogenic cultures, these interactions shape induction outcomes, demonstrating ecological relevance. We extent these findings to infection contexts, showing that arbitrium signalling influences outcomes in cells harbouring prophages with compatible communicating systems. These findings reveal that phages engage in cross-species communication, a trait restricted to more complex life forms, challenging our understanding of how these elements reshape microbial communities.

## INTRODUCTION

Unravelling the foundations of communication within and between species is crucial for comprehending the behaviour of communities and their ecological functions. Communication is not exclusive to advanced animals; even single-celled eukaryotes, bacteria, and bacterial genetic elements like plasmids have developed intricate communication systems.^1–3^ Recently, the concept of inter-cellular communication has been expanded to include viruses, with studies revealing that viruses which infect bacteria (bacteriophages) can make collective life-cycle decisions through small-molecule signalling.^4,5^ Specifically, temperate *Bacillus* phages utilise a small-molecule communication system called arbitrium to regulate the choice between lysis and lysogeny.^5^ This important discovery demonstrated that viruses can behave as complex social entities. Given the significant role bacteriophages (phages) play in bacterial evolution,^6^ understanding these social interactions is essential for grasping the emergence of new bacterial strains.

The arbitrium system consists of three key components: AimP, the arbitrium communication peptide; AimR, the arbitrium receptor, which functions as an anti-terminator protein inhibited by AimP; and *aimX*, a negative regulator of lysogeny.^5,7^ Arbitrum was described in SPβ phages,^5^ but how *aimX* operates in these phages can differ. In some phages, *aimX* encodes a small peptide (AimX) that promotes the lytic cycle of the phage by binding to various partners, including the host’s MazF, thereby inactivating its function as a cellular toxin,^7,8^ while others encode a non-coding RNA in that position whose mechanism is still unknown. ^5,9^At the onset of infection, when the number of infective phage particles is low relative to the number of bacterial cells, the phage begins infecting the population and producing AimP. AimP is synthesised as a 60-80 amino acid (aa) pro-peptide, which, through a proteolytic secretion-internalization process, generates the mature intracellular 6-10 aa active pheromone. Since the extra- and intra-cellular levels of AimP are low at the beginning of the infection, AimR’s function is not inhibited, allowing for the expression of AimX, which promotes the phage’s lytic cycle.^5,10^ However, after several rounds of infection, AimP accumulates extra- and intra-cellularly, which occurs alongside a reduction in the bacterial population, binding to AimR and inhibiting its function. In the absence of AimX, the lysogenic cycle of the phages - characterised by integration into the bacterial chromosome - is favoured, thereby preserving both the bacterial population and the phage within the environment.^5,10^ The arbitrium system not only plays a role during phage infection but is also essential for the proper induction of the integrated prophage.^11,12^

Recent studies have uncovered the existence of thousands of arbitrium-like systems not only in phages, but also in other mobile genetic elements (MGEs), such as conjugative plasmids, and in their bacterial hosts.^9^ Phylogenetic analyses have clustered arbitrium systems into 10 distinct clades, based on homology of the arbitrium AimR proteins.^9^ Each arbitrium system encodes a unique set of mature AimP peptides, varying in size and sequence, and specific to its respective AimR clade. The significant diversity of arbitrium variants and their distinct signals suggests that these systems have evolved to prevent crosstalk among them, and this is the current dogma in the field.^5,9^ This strategy may allow phages to make decisions independently of other phages or MGEs. However, we should not discard the idea that by “listening” to signals from unrelated phages through arbitrium systems, phages could make collective decisions that benefit them as a group. If true, communication between unrelated phages would introduce a new research area with unknown implications for our understanding of bacterial ecology and evolution. Our structural, biochemical, genetic and ecological approaches will demonstrate here that cross-communication via arbitrium systems between unrelated phages does exist and has a profound impact on how phages behave collectively.

## RESULTS

### Exploring cross-talk between arbitrium systems

Although arbitrium systems have been classified into 10 clades, only few systems from clade 2 of the SPβ phage family have been thoroughly characterised to date. These are the arbitrium systems present in phages Spbeta, Phi3T and, to a lesser extent, vB_BsuS-Goe11 (hereafter referred to as Goe11).^11,13–17^ Since the mature AimP in clade 2 is a 6 aa peptide, more than 60 million (20^6^) possible signaling peptides can be generated, easily allowing phage isolation by using different peptides. We carried out a deep search analysis of clade 2 systems identifying more than 286 different AimRs (Table S1). Notably, most mature AimP peptides (73%) associated to these AimRs in this clade share a highly conserved C-terminal “RGA” (Arg-Gly-Ala) motif and the predominant presence of Gly (52 %) or Ser (34 %) at position 1 (Figure 1A; Table S1 and S2), suggesting that the specificity of AimP binding to cognate AimR receptors is mainly determined by two amino acids in position 2 and 3. This fact reduces the number of possible different peptides to the hundreds, a number that should still allow for phages of this clade to evolve in favor of isolating their arbitrium signals away from other species. However, we only found 27 different mature AimPs in this clade (Table S1 and S2). This surprisingly low number strongly suggests that there is evolutionary pressure that limits the number of peptides utilized by the phages. One possible reason could be to facilitate cross-communication between different Arbitrium systems present in different phages. Therefore, to investigate the potential for cross-communication (cross-talk) between arbitrium systems, we initially selected the well characterised Spbeta, Phi3T, and Goe11 phages. The mature AimP peptides for the 3 selected phages are GMPRGA (AimP^Spb^), SAIRGA (AimP^Phi3T^), and GIVRGA (AimP^Goe11^) respectively (Figure 1B). The divergence in these peptides is mirrored by the divergence in their respective AimR receptors (Figure 1C).

**Figure 1.**
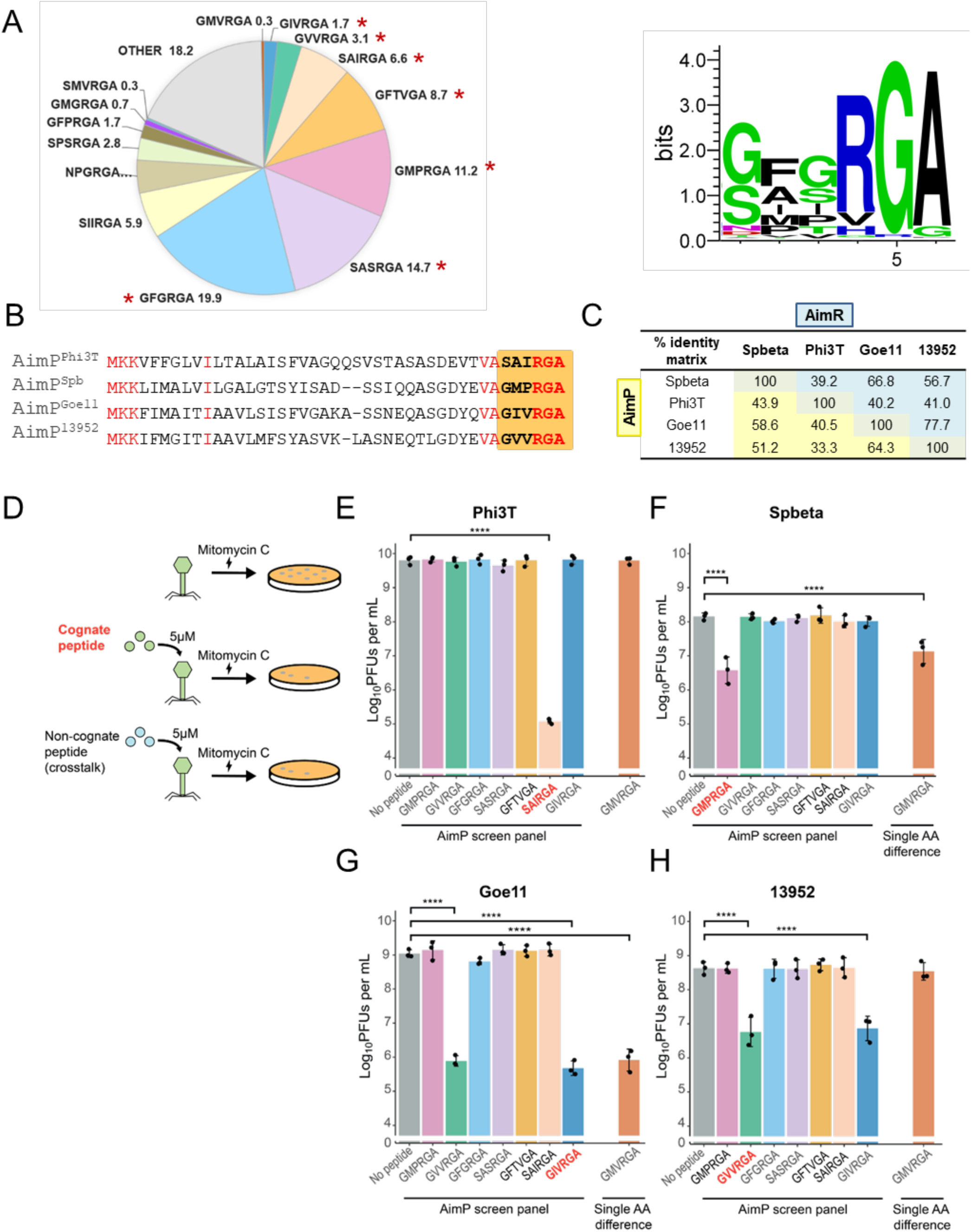
Exploring cross-talk between arbitrium systems. (A) Distribution of AimP sequences in clade 2 (left) and aminoacid profile (Weblogo^43)^ obtained from these sequences (right). Peptides marked with an asterisk are those assayed in subsequent experiments. (B) Alignment of AimP sequences from Spbeta, Phi3T, Goe11, 13952 and V3I1 phages. Conserved residues are painted in red. Mature peptide, corresponding to the last six amino acids is highlighted in orange. (C) Table showing the percent identity matrix calculated with ClustalW^44^ for AimR (shadowed in blue) and AimP (shadowed in yellow) from phages evaluated for crosstalk. (D) Schematic of the protocol for screening potential Arbitrium-mediated crosstalk. A panel of seven synthetic peptides, representative of Clade II arbitrium signals, was exogenously added to phage lysogens harboring different Arbitrium systems. An additional peptide, GMVRGA, which differs in one amino acid from GIVRGA, GVVRGA and GMPRGA, was added to test the extent to which a single amino change in aimP sequence would enable crosstalk (Refer to *Single amino acid changes in AimP peptides can limit cross-talk between arbitrium systems*). These lysogens were then induced by mitomycin C, and the infective phage particles were quantified by conducting plaque assays against *B. subtilis* Δ6s recipient cells (STAR Methods). Columns show log_10_ plaque-forming units (PFUs) mL^−1^ for inductions of: (E) Phi3T, (F) Spbeta, (G) Goe11 and (H) 13952 lysogens. AimP peptides are ordered in decreasing prevalence, based on bioinformatic analysis. A reduction in log_10_ PFUs mL^−1^ relative to the no peptide control indicates the phage responds to the AimP peptide. Cognate peptides for each phage are bolded in red. Columns show the mean of biological triplicates, with standard deviation (SD) as error bars. One-way ANOVAs and post-hoc Dunnett’s tests were conducted on log_10_ transformed data to compare differences, where the culture without exogenous peptide acted as control. P-values are indicated in asterisks above the plot (p < 0.0001, ****). Non-significant p-values were not indicated on the graph.

The exogenous addition of the mature cognate AimP peptide to a lysogenic culture blocks mitomycin C (MC) induction of the resident prophage.^11^ Thus, we hypothesised that testing the ability of synthetically added non-cognate peptides to prevent prophage induction could provide insights into their binding to various AimR proteins (Figure 1D). Specifically, we evaluated a collection of AimP peptides for their capacity to interfere with MC-induced activation of Spbeta, Phi3T, or Goe11 prophages. This collection included AimP^Spb^, AimP^Phi3T^, and AimP^Goe11^, as well as peptides SASRGA, GFTVGA, GFGRGA, and GVVRGA, which are representative clade 2 arbitrium peptides found in other phages.^9^ The AimP peptides studied represent common variants in clade 2, and these seven peptides account for communication in more than 66% of clade 2 phages (Figure 1A;Table S2).

Consistent with the prevailing view that arbitrium systems do not cross-talk, induction of resident prophages was significantly reduced in the presence of their cognate peptides, but not in the presence of the non-cognate peptides (Figure 1E-G). However, an unexpected exception was observed with GVVRGA, which effectively blocked Goe11 induction at levels comparable to AimP^Goe11^ (Figure 1G). This peptide is produced for instance by a prophage present in *Bacillus amyloliquefaciens* strain ATCC13952 (hereafter 13952). Therefore, we included phage 13952 in our studies. In the SPβ family this prophage belongs to the *eta* cluster, unlike Spbeta, Phi3T, and Goe11, which belong to the SPβ cluster.^18,19^ The inclusion of 13952 allowed us to not just explore whether cross-communication exists, but also to investigate if it could occur between phages with the capacity to infect different bacterial species. Since *B. amyloliquefaciens* strain ATCC13952 harboured more than 1 prophage, we generated lysogens of the arbitirum-encoding phage whose AimP is GVVRGA in the same Δ6s *B. subtilis* background strain we used for Spbeta, Phi3T, and Goe11. Note that this phage also can infect this species. Importantly, when testing the ability of the AimPcollection to block 13952 induction, we found that both its cognate GVVRGA peptide (AimP^13952^) and AimP^Goe11^ interfered with MC-induced prophage activation (Figure 1H). This represents a clear example of what we define as symmetric cross-talk, where an AimP peptide produced by one phage can be recognised by an AimR receptor encoded by another phage, and *vice versa*. These results potentially challenge the current assumption that cross-talk does not occur in nature, suggesting that cross-communication between arbitrium systems may indeed be a natural phenomenon.

### Natural AimP peptide levels and their role in prophage induction interference

The previous experiments analysing prophage induction raised an important question: how much AimP peptide is produced in natural conditions? In our earlier experiments, we observed interference in prophage induction by non-cognate peptides using a high concentration (5 µM) of exogenous synthetic peptide (Figure 1E-H). However, the natural concentration of AimP peptides in the supernatant, either after phage infection or during the growth of lysogenic cells, remains unknown. Thus, it was unclear whether the experimental concentrations used were physiologically relevant.

To investigate the effect of AimP peptides produced under natural conditions, we first generated mutant *aimP* phages by deleting the codons encoding the last six amino acids of *aimP* (Δ*aimP^6AA^*), which correspond to the mature peptide (Figure 2A). ^8,11^ Next, we adapted the original protocol used to discover the arbitrium system,^5^ where we collected supernatants from Δ6s *B. subtilis* cultures infected with either the wild type (WT) versions of the Goe11, 13952, Spbeta or Phi3T prophages, or their Δ*aimP^6AA^* mutants. These infected cultures were filtered to remove bacteria and phages, and mixed (50:50) with lysogens carrying the Δ*aimP^6AA^* mutants. The ability of these mixtures to interfere with MC-induced prophage activation was then tested (Figure 2A). Importantly, all supernatants obtained from Δ6s *B. subtilis* strains infected with wild-type phages blocked induction of their cognate prophages, while the supernatants obtained from the infections of Δ*aimP^6AA^* mutants did not affect prophage induction (Figures 2B-2E). Moreover, the supernatants from infections by Goe11 or 13952 also interfered with the induction of each other’s prophages, but not to the Phi3T or Spbeta prophage, strongly supporting the occurrence of cross-talk under natural conditions.

**Figure 2.**
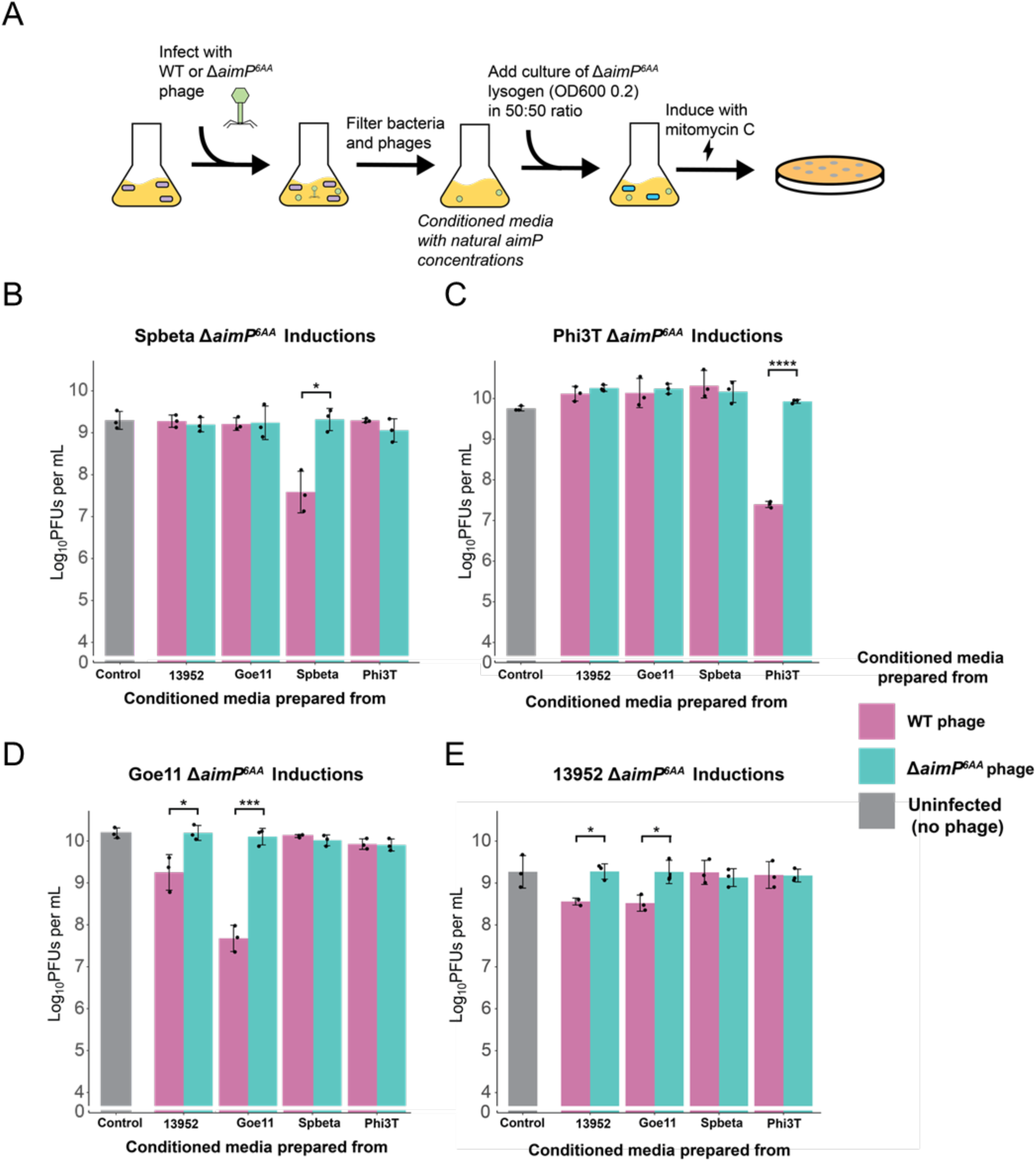
Supernatants of lysogenic cultures containing naturally secreted AimP peptides were sufficient to interfere with MC-induced prophage activation. (A) Schematic showing how conditioned media was prepared and how subsequent induction assays were conducted. Cultures of *B. subtilis* Δ6s were infected with 13952, Goe11, Spbeta, and Phi3T phages, followed by filtration of bacteria and phage particles. The resultant conditioned media containing natural concentrations of arbitirum peptides were mixed with lysogenic phage cultures in equal ratios, followed by MC induction. The infective phage particles were then quantified by conducting plaque assays against *B. subtilis* Δ6s recipient cells. Columns show titers of phage particles in log_10_ PFUs mL^−1^ following MC induction of: (B) Spbeta Δ*aimP*^6AA^, (C) Phi3T Δ*aimP*^6AA^, (D) Goe11 Δ*aimP*^6AA^, or (E) 13952 Δ*aimP*^6AA^ lysogens after mixing with conditioned media. The conditioned media were prepared with the infection of WT or Δ*aimP*^6AA^ phages, indicated by the color of the columns. A reduction in induction titer by conditioned media prepared with WT compared with media prepared with Δ*aimP*^6AA^ phage infection indicates that the interference in induction was due to the naturally secreted AimP peptide. Uninfected control media were prepared, to which no phage was added. Columns show the mean of biological triplicates, with standard deviation (SD) as error bars. Student’s t-tests were conducted on log_10_ transformed data to compare differences in induction titers between conditioned media prepared with WT versus Δ*aimP*^6AA^ phages. P-values are indicated in asterisks above the plot (p < 0.05, *; p < 0.01, **; p < 0.001, ***; p < 0.0001, ****). Non-significant p-values were not indicated on the grap

### *In vitro* binding analysis of cognate and non-cognate peptides to AimR receptors

To gain mechanistic insights into the previous results, we evaluated the ability of cognate and non-cognate peptides, which demonstrated MC-induction interference in earlier experiments, to bind to both their cognate and non-cognate AimR receptors. We hypothesised that, for cross-talk to occur *in vivo*, the affinity of an AimR receptor to a non-cognate peptide should be nearly identical to that observed for its cognate peptide. To test this, we conducted thermofluor assays, a proven method for detecting pheromone binding to AimR and other receptors of the RRNPPA family.^16,17,21,42^[NO_PRINTED_FORM] In these assays, the stability of AimR proteins is assessed in the presence of various AimP peptides. Stronger AimR-AimP binding correlates with a higher temperature required to denature AimR. We produced AimR receptors from Goe11 (AimR^Goe11^), 13952 (AimR^13952^) and from Phi3T (AimR^Phi3T^, used as a negative control), and then measured their melting temperatures (Tm) in the presence of their respective cognate peptides: AimP^Goe11^, AimP^13952^, and AimP^Phi3T^. The results were consistent with previous findings, showing that for both AimR^Goe11^ and AimR^13952^, their cognate and non-cognate peptides (AimP^Goe11^ and AimP^13952^) induce a nearly identical ΔTm, while AimP^Phi3T^ produces significantly lower stabilization (Figure 3A). In contrast, AimR^Phi3T^ exhibited only a strong ΔTm for its cognate peptide compared to the non-cognate ones, explaining why the presence of these non-cognate peptides did not interfere with Phi3T induction.

**Figure 3.**
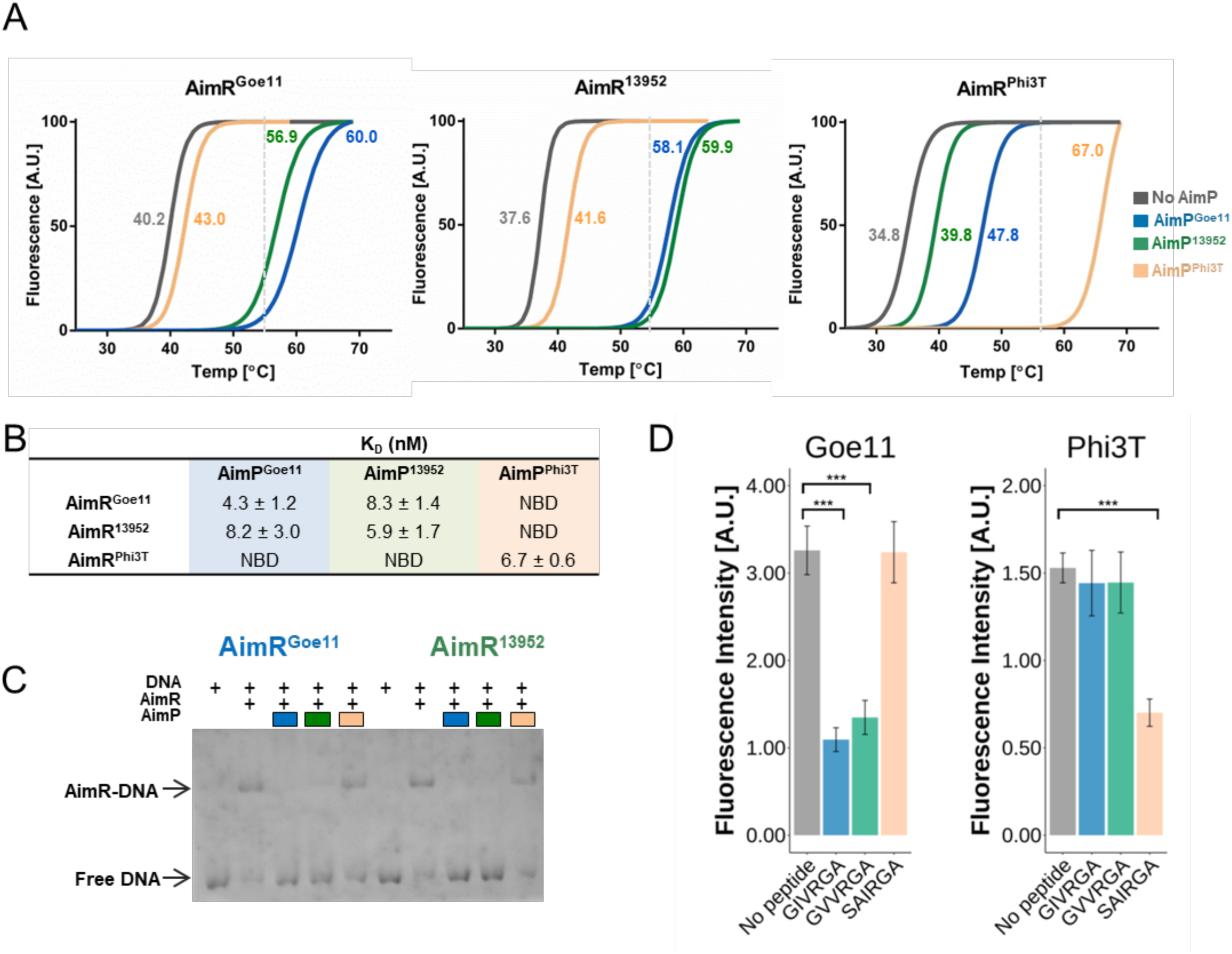
*In vitro* assays of AimR-AimP interactions. (A) Thermofluor shift assay for AimR^Goe11^, AimR^13952^ and AimR^Phi3T^. The vertical lines show the value from which the increase in Tm is greater than 75% of the value reached with the cognate peptide for each AimR. Calculated Tm values in °C are presented next to each curve. (B) K_D_ values calculated by ITC for AimR-AimP binding. NBD denotes “No Binding Detected” at AimP concentrations of 1 mM. (C) EMSA analysis shows that both AimP^Goe11^ and AimP^13952^ peptides induce DNA release in both AimR^Goe11^ and AimR^13952^ while AimP^Phi3T^ has no effect. (D) Fluorescence intensity values after four hours of culture in the presence of the different peptides (0,1 µM). Free AimR allows the expression of sfGFP, but binding of the peptide avoids the effect expression of the reporter.

To confirm these findings and quantify the affinities, we performed isothermal titration calorimetry (ITC) analysis for those AimPs that showed strong stabilization (at least 75% of Tm produced by the cognate peptide). In agreement with the thermofluor results, ITC assays demonstrated nanomolar-range affinities for all receptor-stabilising AimPs, with lower KD values corresponding to AimPs that provided greater stabilisation in thermofluor assays. The stabilisation of AimR^Goe11^, AimR^13952^ and AimR^Phi3T^ by their respective cognate pheromones was confirmed by affinity constants ranging from 4.3 to 6.7 nM (Figure 3B and Figure S1). The affinities of AimRs for their non-cognate peptides (AimR^Goe11^-AimP^13952^ and AimR^13952^-AimP^Goe11^) was only slightly lower than for their cognate peptides (8.2 and 8.3 nM) (Figure 3B and Figure S1).

Next, we validated the functionality of non-cognate peptides using electrophoretic mobility shift assays (EMSA). In these experiments, the formation of the AimP-AimR complex inhibits AimR from binding to its DNA operator sequence.^16^ Notably, all peptides that showed significant binding to the AimRs also strongly affected AimR function (Figure 3C), supporting the hypothesis that certain non-cognate peptides can modulate AimR activity in unrelated phages. Finally, to demonstrate that the observed effect with the non-cognate peptides was due to their ability to bind to AimR and inhibit its function, we tested the anti-terminator activity of AimR using transcriptional reporters. These reporters were constructed by fusing the regions downstream of the transcriptional terminator, whose function is inhibited by AimR, to sfGFP, and the expression of the reporter was measured for AimR^Goe11^ and AimR^Phi3T^ in the presence of the different peptides. In support of the previous results, the cognate AimP^Goe11^ and the non-cognate AimP^13952^ peptides were able to disrupt AimR^Goe11^ anti-termination function, blocking GFP expression, while for AimR^Phi3T^, only the cognate AimP^Phi3T^ was able to do so (Figure 3D).

### Molecular insights into arbitrium cross-talk

To further understand the molecular basis of cross-communication in arbitrium systems, we characterized the structure of the AimR receptors showing cross-talk in the presence of their cognate and non-cognate AimPs. We have already solved the 3D structures of different AimR receptors, including AimR^Goe11^, in their apo and cognate AimP-bound conformations.^16,17^ Therefore, we next solved the structure of AimR^13952^ alone and in complex with AimP^13952^, which was done at 2.3 and 3.1 Å resolution, respectively (Table S3). Both structures showed the presence of two molecules in the asymmetric unit forming a dimer, confirmed in solution by SEC-MALS (Figure S2). Residues 1-44, corresponding to part of the DNA Binding Domain (DBD), exhibited high mobility, preventing them from being traced in the final model of both structures. The folding of the remaining part of the protein (residues 45-386) is similar to that found in AimR^Goe11^, and each protomer includes nine degenerated tetratricopeptide repeats (TPRs) arranged in two subdomains, TPR_Nter (TPR1-6) and TPR_Cter (TPR7-9), connected by a linker (Figure 4A, 4B and Figure S3). The structure of AimR^13952^ in complex with AimP^13952^ (AimR-AimP^13952^) confirms the closure movement of both TPR subdomains, previously observed in the AimP-bound structures of AimR^SPβ^ and AimR^Goe11^, which compacts each subunit in the dimer (Figure 4C).^16,17^

**Figure 4.**
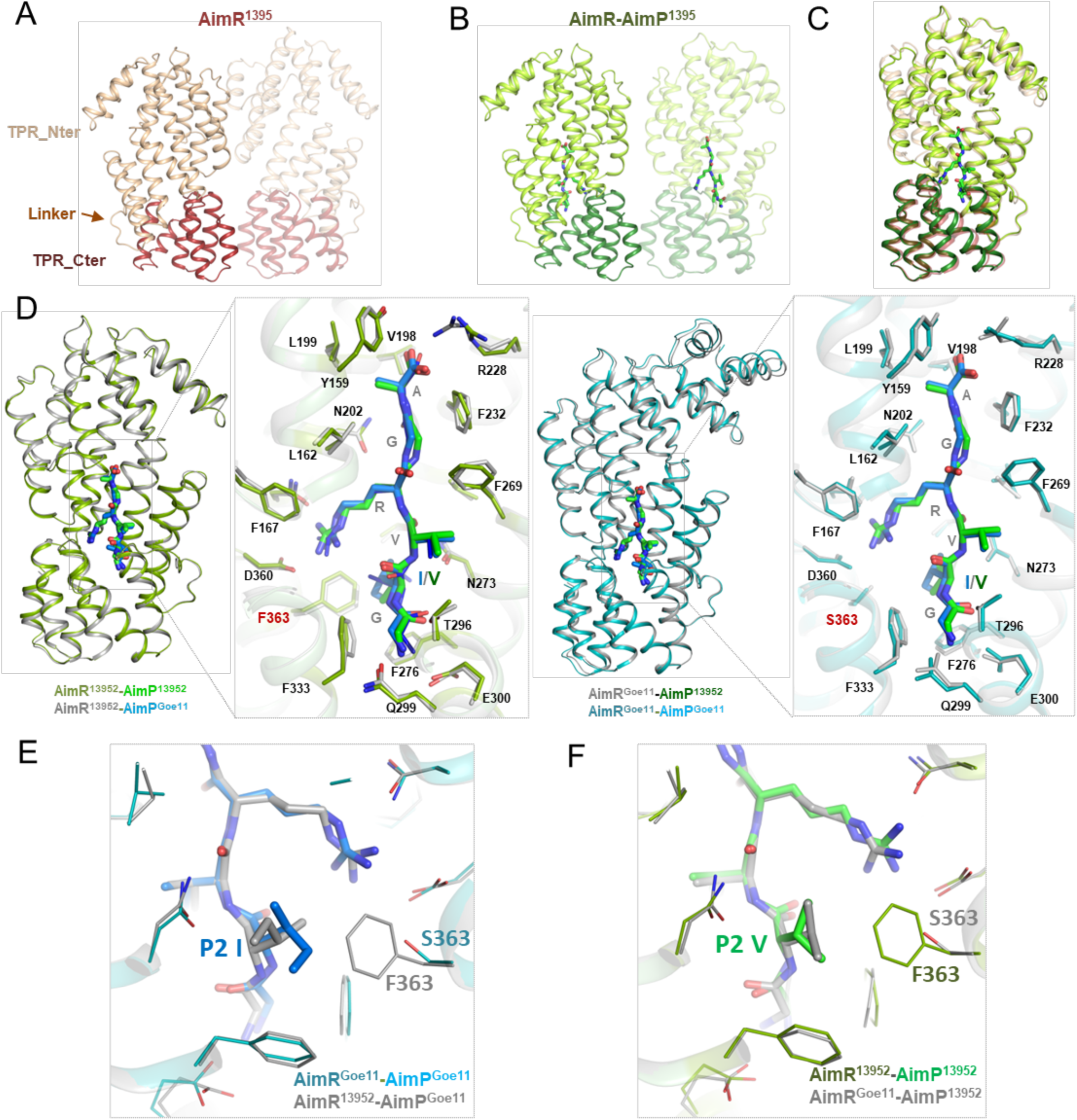
Structures of AimR^13952^ and AimR-AimP complexes. (A) Left, overall structure of AimR^13952^ dimer with TPR N-ter domain painted in beige and TPR C-ter domain painted in red. Linker connecting both domains is marked with an arrow. In the middle, overall structure of AimR-AimP^13952^ complex with AimP showed as green sticks. TPR N-ter and TPR C-ter domains are coloured in light and dark green, respectively. In both figures monomer B is shown in semitransparent cartoon, using the same colours. In the right, superposition of AimR^13952^ and AimR-AimP^13952^ monomers. (B) Superposition of each AimR with its cognate and non-cognate AimPs, shown as sticks, and colored in blue for AimP^Goe11^ or green for AimP^13952^. Complexes with canonical peptides are colored in green for AimR-AimP^13952^ or blue for AimR-AimP^Goe11^. Complexes with non-canonical peptides are colored in grey. A close view of peptide binding sites of the superposed AimRs is presented in the boxes. (C) Superposition of AimR13952-AimP13952 (in green) and AimR^Goe11^-AimP^13952^ (in grey). The presence of a Ser at AimR^Goe11^ position 363 allows the accommodation of Val at AimP position 2. (D) Superposition of AimR^Goe11^-AimP^Goe11^ (blue) and AimR^13952^-AimP^Goe11^ (grey). The presence of a Phe at AimR^13952^ position 363 forces Ile at AimP^Goe11^ position 2 to adopt an alternative conformation.

Next, we solved the three-dimensional structure of AimR^Goe11^ and AimR^13952^ in complex with the reciprocal non-cognate peptides (AimP^13952^ and AimP^Goe11^), obtaining the complexes AimR^Goe11^-AimP^13952^ and AimR^13952^-AimP^Goe11^ at 1.9 and 2.6 Å resolution, respectively (Figure S4A and S4B; Table S3). Superposition of the structures of each AimR in complex with their cognate and non-cognate AimP showed that both peptides induce nearly identical conformational changes with RMSDs of 0.51 Å (AimR^13952^) and 0.46 Å (AimR^Goe11^) for the receptor subunit pairwise comparison (Figure S4). In fact, the same structural analysis for the four structures (both AimR receptors in complex with cognate and non-cognate AimPs) confirms the matching conformation (RMSDs of 0.46 to 0.87 Å) in all cases (Figure S4C and S4D). A close view of the interacting AimR residues in the complexes revealed that the AimPs are recognised by 19 residues, of which 14 interact with the side chains of the AimPs, providing specificity by the peptide (Figure 4D; Table S4). All these residues, except one (position 363), are conserved between AimR^Goe11^ and AimR^13952^; the differing residue is a Ser in AimR^Goe11^ and a Phe in AimR^13952^. As might be anticipated, this residue is in the vicinity of the AimP position 2 (P2) (Figure 4D), which is the amino acid by which the peptides AimP^Goe11^ (GIVRGA) and AimP^13952^ (G<u>V</u>VRGA) differ. The change from a Ser (AimR^Goe11^) to a Phe (AimR^13952^), which introduces a much more voluminous and hydrophobic side-chain, reduces the space to accommodate the side chain of the AimP P2 amino acid. This explains why AimP^13952^ has a Val at the P2 position, the side chain of which is smaller than the corresponding Ile of AimP^Goe11^. However, the structure of the AimR^13952^-AimP^Goe11^ complex shows that this reduction in space does not prevent the accommodation of AimP^Goe11^. Instead, the P2 Ile of this AimP^Goe11^ can adopt an alternative conformation, where it has hydrophobic interactions with the AimR^13952^ Phe at position 363 without causing steric problems (Figure 4E). In the case of AimP^13952^, the smaller Val at P2 has no problem accommodating itself in AimR^Goe11^, where there is a much larger space as this receptor has a Ser at position 363 (Figures 4F). Taken together, the structures show at the molecular level how the AimR receptors have made specific changes to maintain high specificity for their AimP without reducing affinities for those of other systems, with which they may engage in cross-talk.

### *In vivo* evidence of arbitrium cross-talk

To further validate the findings that crosstalk among arbitrium systems occurs, we generated chimeric phages expressing non-cognate AimP peptides. These chimeric phages retain the original *aimP* promoter present in the phage, with only the DNA region encoding the last six amino acids of AimP replaced (Figure 5A). We constructed the chimeric derivatives of Goe11, 13952, Spbeta, and Phi3T harboring the 6 amino acid peptides of each other. We hypothesised that when there is a natural interaction between the expressed AimP and AimR, the titre obtained after MC induction of these prophages would be similar to that obtained after induction of the WT phage, and much lower than that obtained after induction of either the Δ*aimP^6AA^* phage or a chimeric phage expressing a non-cognate AimP which is unable to interact with AimR. As predicted, induction of the Goe11 Δ*aimP^6AA^* mutant and the Goe11 chimeric phages expressing AimP^Phi3T^ and AimP^Spb^ produced 1000 times more infective particles than the WT phage (Figure 5B). However, the chimeric Goe11 phage expressing AimP^13952^ produced titres identical to the WT Goe11 phage. Similarly, the chimeric 13952 phage expressing AimP^Goe11^ suppressed infective particles to WT levels, while other chimeric derivatives induced to the same level as the Δ*aimP^6AA^* mutant (Figure 5C). Meanwhile, inductions of all Spbeta and Phi3T chimeric derivatives produced identical numbers of infective particles to the Δ*aimP^6AA^* phage (Figure 5D, 5E). Thus, these results confirm that naturally produced non-cognate peptides are sufficient to inhibit AimR^Goe11^ and AimR^13952^ function.

**Figure 5.**
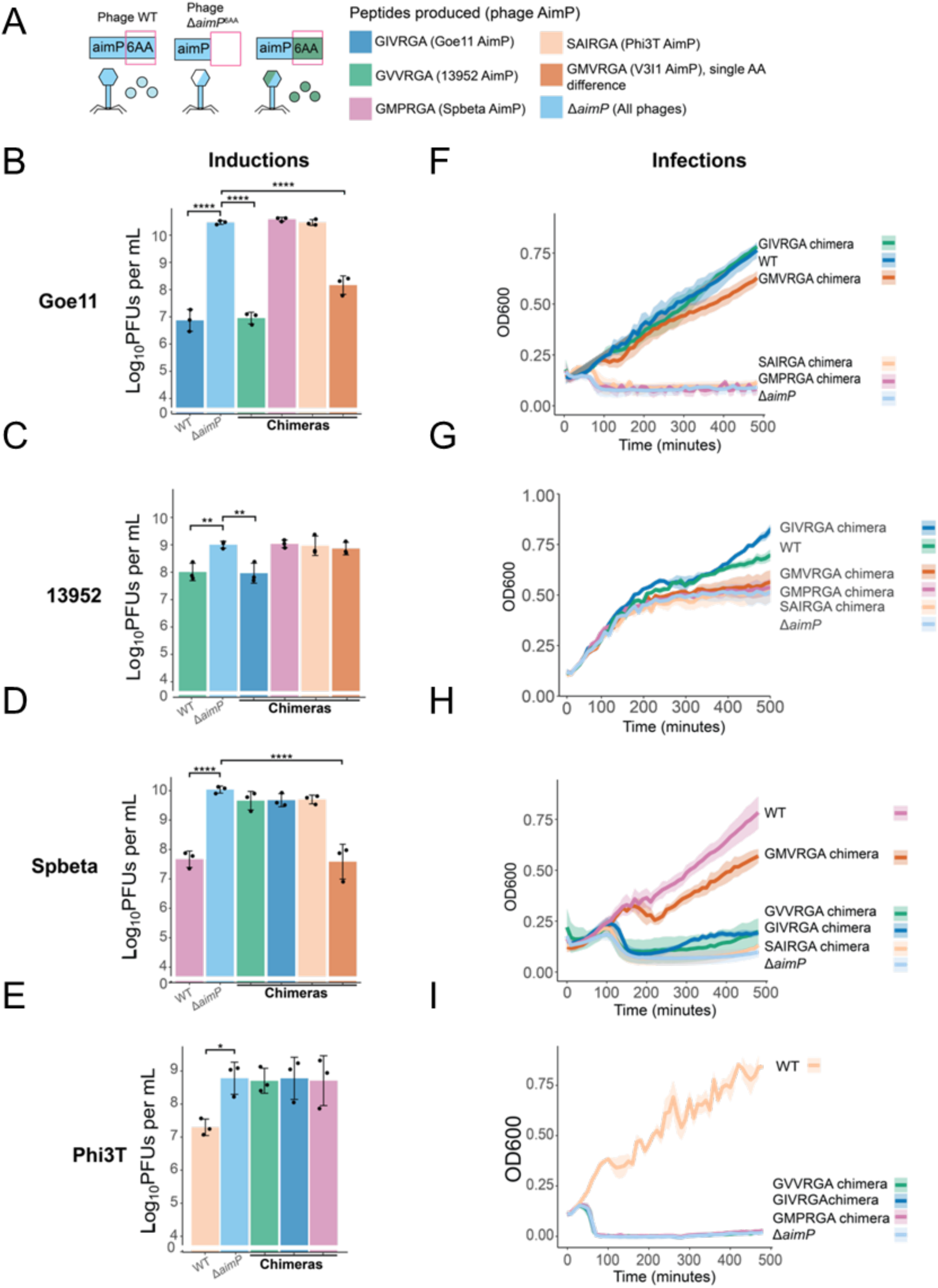
Chimeric phages expressing mature non-cognate AimP peptides behaved similarly to WT phages in both induction and infection. (A) Chimeric AimP phages were cloned, in which the last six amino acids of AimP encoding the mature peptide were swapped for non-cognate sequences. These derivatives were induced, along with their WT and Δ*aimP*^6AA^ counterparts, and quantified with plaque assays. The columns show titers in log_10_ PFUs mL^−1^. Lysates of the phages derivates were also added to cultures of *B. subtilis* Δ6s at MOI 0.1, the survival of which was then tracked on growth curves. Color scheme shows the different peptides produced by the phages and their chimeras. For phages Goe11, 13952, and Spbeta, an additional aimP derivative was cloned, such that the AimP belonging to phage V3I1 (GMVRGA) was produced, which differs in one amino acid to the aforementioned phages (Refer to *Single amino acid changes in AimP peptides can limit cross-talk between arbitrium systems*). Induction titers of (B) Goe11, (C) 13952, (D) Spbeta (E), and Phi3T chimeras. Growth curves of (F) Goe11, (G) 13952, (H) Spbeta (I), and Phi3T chimeras. Columns in induction bar graphs show the mean of biological triplicates, with standard deviation (SD) as error bars. Shadows in growth curves indicate standard error. One-way ANOVAs and post-hoc Dunnett’s tests were conducted on log_10_ transformed data to compare differences in induction titers with Δ*aimP*^6AA^ as control. P-values are indicated in asterisks (p < 0.05, *; p < 0.01, **; p < 0.001, ***; p < 0.0001, ****). Non-significant p-values were not indicated here

Next, we evaluated the behaviour of the WT, Δ*aimP^6AA^*mutant, and chimeric Goe11 phages during infection. This was the original scenario used to identify the arbitrium system,^5^ and here, the interaction between AimP and AimR will promote the survival of the culture after infection by promoting lysogeny. Infection with WT Goe11 or the chimeric Goe11 phage expressing AimP^13952^ displayed significantly increased cell survival compared with infection with the Δ*aimP^6AA^* mutant or the chimeric phage expressing other non-cognate AimPs (Fig. 5F). Initial cell death due to lytic activity was followed by lysogeny-mediated culture protection, indicating clear arbitrium-mediated control induced by the cognate peptide and AimP^13952^ (Fig. 5F). Similarly, infection of WT 13952 phages and chimeric 13952 derivative expressing AimP^Goe11^ had increased survival compared to the Δ*aimP^6AA^* mutant and other derivatives (Fig. 5G). However, in the case of Phi3T and Spbeta, only the cognate AimPs promote culture survival (Figure 5H and 5I). Taken together, these results demonstrate that the natural production of AimP peptides, either during infection or by lysogenic cells, is sufficient to mediate symmetrical cross-talk between arbitrium systems of Goe11 and 13952, confirming that this process can exist in nature.

### Single amino acid changes in AimP peptides can limit cross-talk between arbitrium systems

The observation that AimP peptides produced by the Goe11 and 13952 phages differed by only one position raised the question of whether the observed cross-talk was specifically selected or would occur universally when AimP peptides differ by a single amino acid. To address this, we investigated the peptide produced by a phage from *Bacillus atrophaeus* V3I1. The AimP peptide encoded by this phage (AimP^V3I1^) has the sequence GMVRGA, which differ from the peptides encoded by Goe11 (GIVRGA), 13952 (GVVRGA), and Spbeta (GMPRGA) by one amino acid in P2 or P3. This allowed us to test this peptide with three different phages. Notably, when AimP^V3I1^ was added exogenously, it hindered MC-induced prophage activation of Goe11 and Spbeta, therefore representing another example of potential inter species cross-communication (Figure 1F-H). Importantly, it did not block induction of 13952.

To confirm whether this interference can also be observed at natural concentrations, we generated chimeric Goe11, 13952, and Spbeta derivatives encoding AimP^V3I1^ (Figure 5). Notably, Spbeta derivatives producing this peptide induced to WT levels (Figure 5D); for Goe11, the chimeric derivative had lower phage titres than Δ*aimP^6AA^* and the blocking was close to the extent of WT or AimP^13952^ (Figure 5B). For 13952, the chimeric phage producing AimP^V3I1^ had the same titres as its Δ*aimP^6AA^* mutant (Figure 5C), aligning with our observations with the synthetic peptide screens. Similarly, when we tested infections with these phage derivatives, we found that Goe11 and Spbeta derivatives expressing AimP^V3I1^ displayed increased cell survival than the Δ*aimP^6AA^* phage, but infection with the 13952 AimP^V3I1^ derivative displayed cell survival similar to that of Δ*aimP^6AA^* (Figure 5F-H).

These *in vivo* results correlate with our previous structural analysis of peptide specificity. AimP^V3I1^ differs with AimP^Goe11^ and AimP^13295^ only at the P2 position, introducing a longer Met residue (vs Ile/Val). The increment in size is not a problem for Goe11, whose AimR faces this AimP position with a small Ser (363), but the presence of a bulky Phe for 13952 would generate steric problems that prevent AimP^V3I1^ binding (Figure 4E and 4F). Therefore, changes in the peptide’s position can be compensated with minimal changes in AimR’s recognition residues, thereby allowing for different degrees of binding promiscuity. These findings suggest that even a single amino acid substitution in the AimP peptide can modulate cross-talk between unrelated phages to varying levels of sensitivity by changing punctual positions in AimR recognition residues, ranging from-mediated control at all. In this way, it is to be expected that some phages have opted for a higher level of promiscuity while others severely restrict their communication. Our results also highlight that crosstalk may occur frequently in nature, as we report here other examples (Goe11 and V3I1; SPBeta and V3I1) where these interactions take place. In summary, our results indicate that the observed cross-talk is a bona fide interaction between specific AimP peptides and AimR receptors from unrelated phages, rather than a universal feature of closely related AimP peptides. This highlights the specificity of the arbitrium system in mediating communication among phages.

### Arbitrium cross-talk shapes phage interactions in complex ecological contexts

The previous results suggested that in mixed populations, where different lysogenic strains coexist, the arbitrium-mediated cross-talk interactions between phages could play a crucial role in shaping their dynamics. To evaluate this idea, and to provide additional evidence of the role this process has in a more natural scenario, we mixed lysogenic cells carrying either the WT or the Goe11 *ΔaimP^6AA^* mutant prophage with lysogenic cells harbouring either the WT or *ΔaimP^6AA^* mutant of the 13952 prophage in a 50:50 ratio. The mixed cultures were then induced with MC and the infective particles produced by each phage were quantified separately (Figure 6A). In all combinations tested, the presence of at least one prophage encoding its cognate *aimP* was sufficient to reduce the titres of both phages after induction. In contrast, induction of the community carrying both prophages mutated in Δ*aimP^6AA^* resulted in significantly higher titres for both phages (Figure 6B and 6C). This indicates that the arbitrium peptides produced by either Goe11 or 13952 are sufficient to inhibit induction of both cognate and non-cognate phages to a similar extent, with no detectable difference based on the source of the peptide. By contrast, in the absence of any AimP peptide, both prophages are strongly induced due to the lack of AimP-mediated repression of prophage activation. These results confirm an important role for cross-talk between arbitrium systems in shaping the ecological interactions and coordinated behaviour of phage communities.^20^

**Figure 6.**
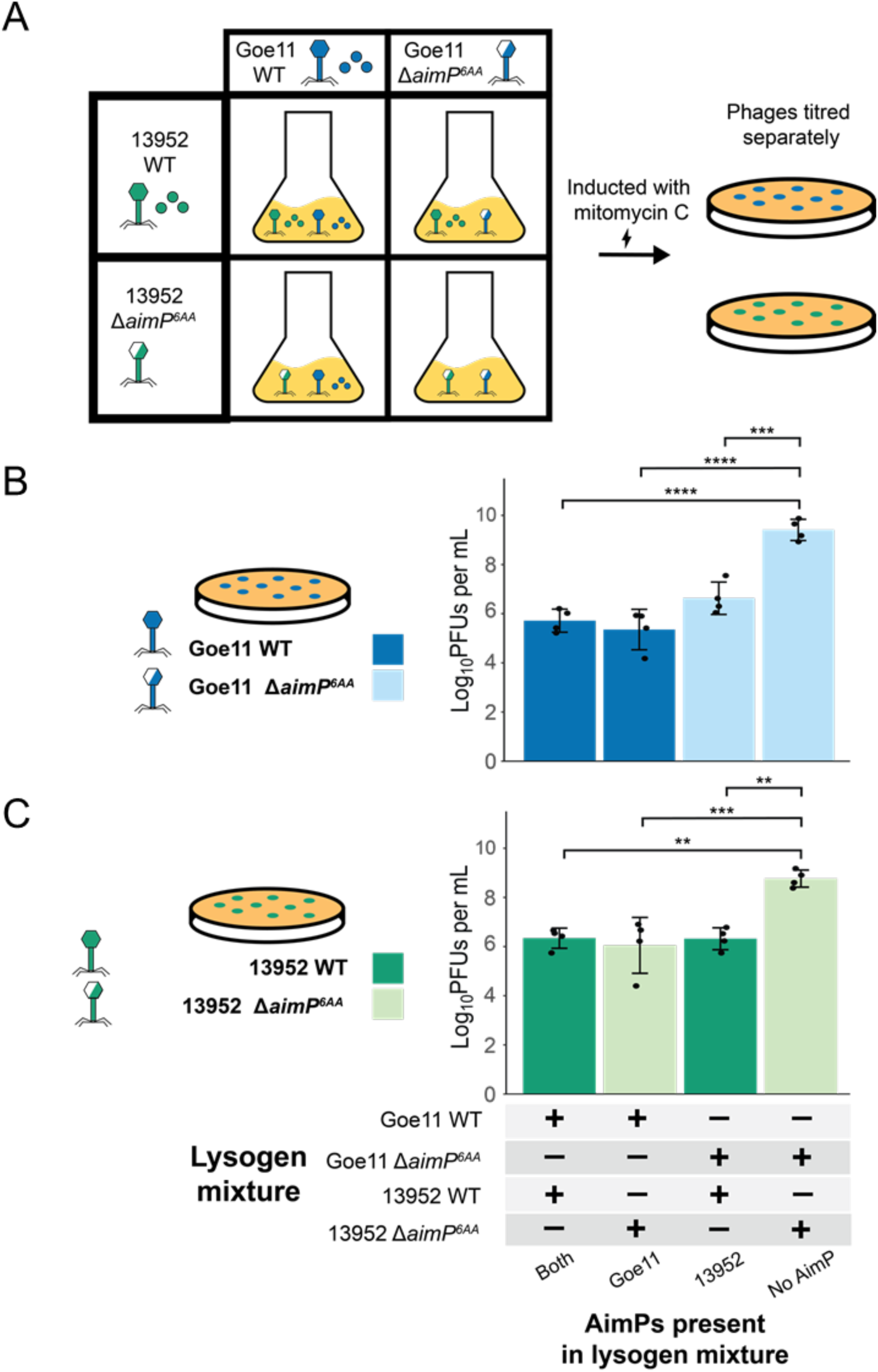
Arbitrium crosstalk shapes phage interactions in mixed populations. (A) Schematic overview of protocol for mixed population inductions. Cultures of WT and Δ*aimP*^6AA^ phages were independently grown to OD600 0.2 and mixed in equal volumes. These mixtures were then MC induced and infective phages particles were quantified separately in plaque assays, using different bacterial recipients, which are represented in the column graphs below. (B) Induction titers of Goe11 phages by mixture; (C) induction titers of ATCC13952 phages by mixture. The columns are vertically aligned such that Goe11 and ATCC titers in the same position on the x axis were part of the same mixture. Plus and minus symbols underneath the columns indicate presence and absence of lysogens in the mixture respectively. The color of columns indicates whether the Goe11 and ATCC13952 added to the mixture was WT or Δ*aimP*^6AA^. The resultant aimP present in the mixture is indicated at the bottom. Columns show the mean of biological triplicates, with standard deviation (SD) as error bars. One-way ANOVAs and post-hoc Tukey’s tests were conducted on log_10_ transformed data to compare differences in induction titers between mixtures. P-values are indicated in asterisks above the plot (p < 0.05, *; p < 0.01, **; p < 0.001, ***; p < 0.0001, ****). Non-significant p-values were not indicated on the graph.

To complete this study, we next analysed whether cross-talk between different phages also impacts phage and bacterial communities during infection. Our previous results with chimeric phages also suggest that during phage infections, arbitrium cross-talk protects the bacterial population by guiding the phage to lysogeny. Hence, we hypothesize that in a natural scenario where a phage infects a population of lysogens containing a cross-talking phage, the lysogen population should impact the lysogeny of the incoming phage, thereby protecting the bacterial population. To test this idea, we grew cultures of lysogens harbouring different derivatives of 13952 (WT, Δ*aimP^6AA^*, chimera encoding AimP^Goe11^, chimera encoding AimP^Phi3T^), and then infected them with WT Goe11 encoding a tetracycline marker (Figure 7A). After 1.5 hours, which corresponds to the time when maximum lysis occurs, the infection mixtures were plated on either LB plates or LB with tetracycline plates to quantify the number of total cells and Goe11 lysogens respectively. The differences in total number of cells surviving the infection between the various recipients were minimal, likely because the survival rate was high in all cases (Figure 7B). Importantly, the number of Goe11 lysogens that formed when infecting lysogens that harboured 19352 WT or the AimP^Goe11^ chimera was significantly higher than when infecting Δ*aimP^6AA^* mutant or AimP^Phi3T^ chimera (Figure 7C), confirming that arbitrium-mediated crosstalk can modulate the outcome of infections by cross-talking phages.

**Figure 7.**
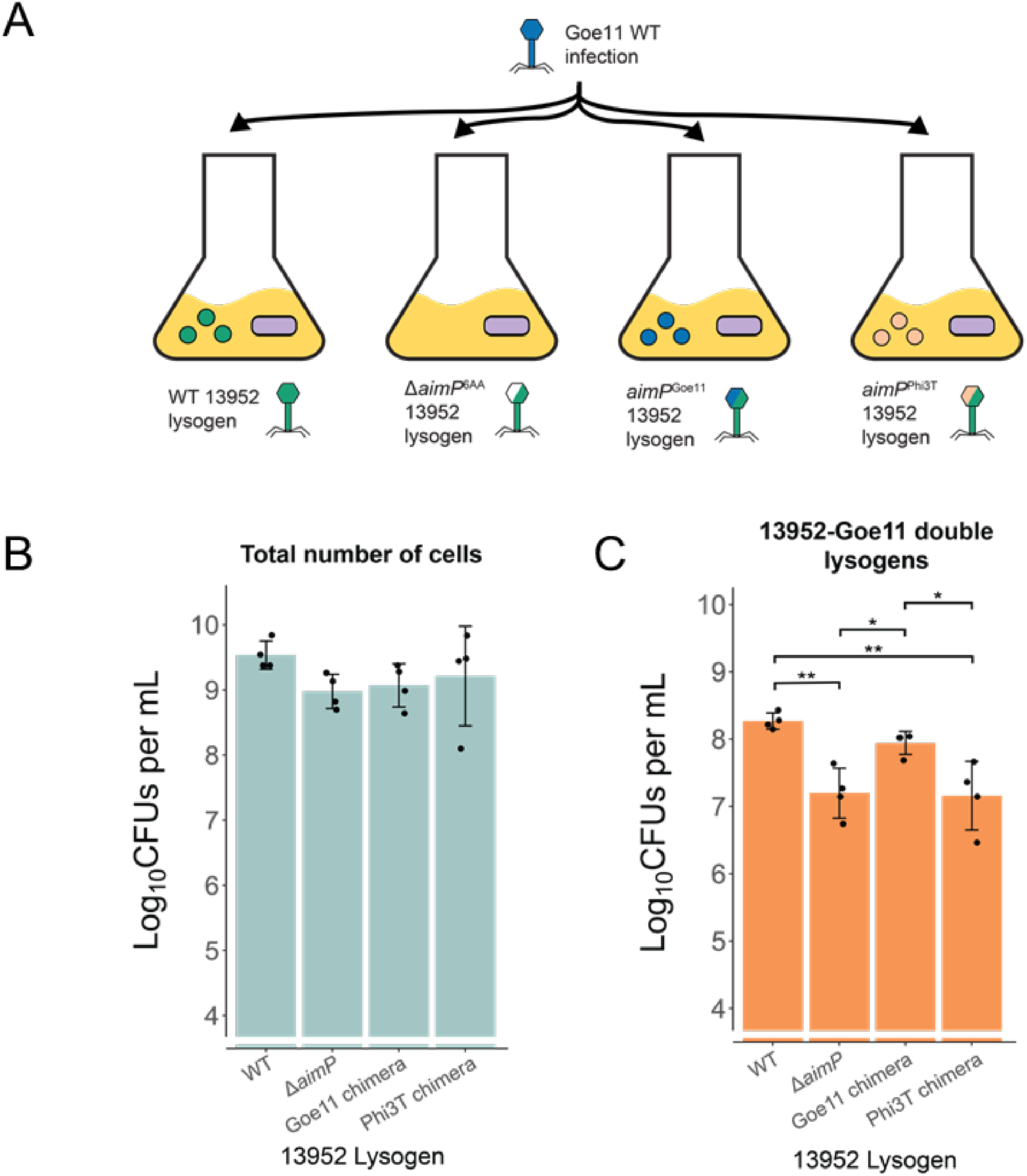
Arbitrium crosstalk shapes infection outcomes of incoming phages. (A) Schematic overview of protocol for infection dynamic experiments. Cultures of lysogens harbouring different derivatives of 13952 phages were grown to OD600 of 0.2, followed by infection with WT Goe11 (MOI 1) encoding a tetracycline marker. After 1.5 hours, the infection mixtures were then plated on LB and tetracycline-containing LB plates to quantify the number of total cells and Goe11 lysogens respectively. (B) Total number of cells and (C) number of Goe11 lysogens for each infection. Columns show the mean of biological triplicates, with standard deviation (SD) as error bars. One-way ANOVAs and post-hoc Tukey’s tests were conducted on log_10_ transformed data to compare differences. P-values are indicated in asterisks (p < 0.05, *; p < 0.01, **). Non-significant p-values were not indicated on the graph.

Overall, these findings confirm the existence of arbitrium-mediated cross-talk and their impact and ecological relevance in nature. Remarkably, in these experiments we analyse phages originating from *B. amyloliquefaciens* (13952) and *B. subtilis* (Goe11), two species that coexist,^20^ indicating that cross-talk can occur among phages from different bacterial species.

## Discussion

Our findings extend the paradigm of phage communication, demonstrating for the first time that arbitrium systems can mediate cross-talk between unrelated phages, which may infect different bacterial species. This challenges the assumption that, by using arbitrium, phages only communicate with their progeny^5^. ^5^ Remarkably, we observe cross-talk between peptides produced by phages that reside in three different *Bacillus* species (*B. subtilis*, *B. amyloliquefaciens and B. atrophaeus)*, including those sharing the same ecological niche,^20^ which highlights the ecological relevance of this phenomenon. Additionally, other MGEs, apart from phages, also encode arbitrium systems,^9^ whose functions and potential interactions remain to be explored. The ability of certain phages to communicate not only with their progeny, but also with unrelated phages or other MGEs, has profound implications for understanding bacterial and phage ecology.^22,23^ By responding to signals from unrelated phages, these viruses may collectively optimise their life cycle decisions, potentially influencing the dynamics of bacterial populations and the spread of mobile genetic elements.^24^ This discovery adds a new layer of complexity to viral social behaviours and opens avenues for investigating the ecological roles of inter-phage communication. Our results may represent just the tip of the iceberg, suggesting a vast, uncharted network of inter-phage and MGE communication systems yet to be uncovered.

The observed cross-talk raises intriguing questions about the evolutionary pressures shaping arbitrium systems. While specificity in AimP-AimR interactions may have evolved to prevent detrimental interference between phages, the fact that the number of distinct peptides found in nature is significantly lower than the number theoretically possible suggests that some phages may have evolved arbitrium systems to promote interaction between unrelated systems. We have confirmed this idea showing that cross-communication could confer selective advantages under certain ecological contexts. For instance, cross-talk might enable phages to sense the density of unrelated viral populations or to coordinate strategies that preserve host populations during co-infections. While it is clear that cross-talk can be detrimental for some phages under certain scenarios - such as when it limits the replication of an incoming phage infecting a lysogen capable of cross-communication - it is equally evident that it confers advantages in other contexts. For example, our experiments demonstrate that it can promote the existence of double lysogens. It is well known that prophages often encode accessory genes, including anti-phage systems^27^. In some scenarios, these double lysogens may survive better than single lysogens, particularly if the prophages carry genes that protect the bacterial population.

Importantly, because cross-talk can be detrimental, phages may have evolved mechanisms to minimize interactions with unrelated phages. Remarkably, a single amino acid change in the pheromone can isolate one system from others. In contrast, our structural results show that AimRs can evolve to recognize both cognate and non-cognate peptides with nearly identical affinity, thereby increasing their promiscuity in communication. These minimal AimR changes can be evolutionarily selected to decide, through pheromone recognition, with which group of phages to communicate or, alternatively, to remain isolated. Therefore, our structural and biochemical analyses revealing the subtle molecular determinants of AimP specificity underscore how minor variations in pheromone-receptor interactions can mediate this phenomenon, balancing specificity with the potential for beneficial cross-communication. We anticipate that the ecological niche and the population of MGEs present should be close determinants in these variations.

Our results also demonstrate that arbitrium-mediated cross-communication can occur at varying sensitivities, where one cross-talking phage (Spbeta) may be more sensitive to the same non-cognate peptide (GMVRGA from V3I1) than another phage (Goe11). It is tempting to speculate that such variance in peptide sensitivity may allow different phages to modulate cross-talk to varying degrees. Some phages may evolve to become highly sensitive to non-cognate peptides with the primary intent on ‘eavesdropping’ the lysis-lysogeny decisions of other phages. In contrast, other phages may utilize crosstalk as ancillary information, thereby making smaller adjustments to lysis-lysogeny decisions that are primarily guided by the cognate peptide. Variations in sensitivity may depend on how much of the biological niche is shared between the phages and bacterial host, how much molecular machinery is shared between the competing phages, or even the presence of other modulating anti-phage systems. This can allow a part of the phage population to adjust its lifecycles and balance towards a prudent infection^24^ that does not deplete the host population.^22^

This study not only challenges existing paradigms but also sets the stage for future research into the ecological and evolutionary roles of viral communication. Investigating the prevalence of cross-talk in natural environments and its impact on microbial ecosystems will be crucial for understanding its broader significance. Additionally, the insights gained here could inform the development of novel biotechnological tools, such as engineered phages with customised communication systems for targeted bacterial control. By harnessing the principles of arbitrium-mediated communication, we may unlock new strategies for combating bacterial pathogens and mitigating antibiotic resistance.

## STAR Methods

### Bacterial strains and reagents in this study

All bacteria used in this study are listed in Table S5 and are derivates of *E. coli* DH5α (NZTtech), *E. coli* strain BL2_Codon plus (DE3) RIL (Agilent) or *B. subtilis* Δ6s. *B. subtilis* Δ6s is a chloramphenicol-sensitive derivative of the *B. subtilis* Δ6 strain (NCBI Ref: NZ_CP015975.1), which lacks mobile genetic elements. Bacteria were either grown on LB Lennox media/agar (*E. coli*), LB Miller media/agar (*B. subtilis*), or Bacillus minimal media (protocol from^25^). Competent *E. coli* cells were purchased from NZTtech or Agilent and competent *B. subtilis* cells were prepared as previously described.^30^ Synthetic AimP peptides were purchased from ThermoFisher Scientific and Proteogenix (>95% purity, desalted). Synthetic oligonucleotides were purchased from Sigma-Aldrich and IDT and listed in Table S5. Sanger sequences were done by Eurofins and Macrogene, and short-read Illumina sequencing was done by Seqcenter.

To isolate the prophage Goe11 from the strain *B. subtilis* strain TS01 (gift from Dr. Robert Hertel^19^) and the Arbitrium-harboring prophage in *Bacillus amyloliquefaciens* ATCC13952, the bacterial strains were first induced with mitomycin C (MC) to obtain lysates of resident prophages. Plaque assays were then conducted to obtain individual phage plaques, which were then stabbed and suspended in phage buffer. PCRs were used to identify suspensions harboring the prophage of interest. Positive suspensions were then replated with further plaque assays and verified with PCRs again twice more to obtain pure phage lysates. The lysate was then spotted onto a lawn of *B. subtilis* Δ6s to obtain phage lysogens.

### Molecular cloning

*B. subtilis* Δ6s was generated by deleting the pks::cmR marker in *B. subtilis* strain Δ6. The deletion was made with the Cre-lox recombination system. Briefly, the *cmR* marker was replaced with a *kanR* cassette flanked by loxP sites. The plasmid pDR244 was then transformed with spectinomycin selection (100 µg/ml) at 30°C to allow Cre-loxP-mediated removal of *kanR*. Transformants were then streaked onto LB agar at 42°C to cure the plasmid. Strains were screened for spectinomycin and chloramphenicol sensitivity, streaked to single colonies, and verified with PCRs.

To quantify 13952 titres in mixed population lysates, a Δ*ggaB* mutant of *B. subtilis* Δ6 was generated. Briefly, triple PCRs were used to amplify the erythromicyn cassette *ermR* with loxP sites and flanking sequences overlapping *ggaB*. The PCR product was then transformed into Δ6 to replace *ggaB*. The *ermR* cassette was then removed by using pDR244 as before.

To quantify Goe11 titres in mixed population lysates, the 13952 master repressor gene, *sroF* (*sroF^13952^*), was cloned into the recipient *B. subtilis* Δ6s. The gene was cloned under an IPTG-inducible promoter, pSpank, using the amyE integrative plasmid pDR110.^26^ Briefly, *sroF^139^*^52^ was amplified by PCR, with BamHI and EcoRV restriction sites introduced as overhangs. The fragment was then ligated with the linearized pDR110, and transformed into *B. subtilis* Δ6s. Strains were screened for spectinomycin resistance (100 µg/ml) at 37°C and loss of amylase activity by starch test. Strains were then streaked to single colonies and verified with PCRs and Sanger sequencing. The same protocol was used to clone the Goe11 master repressor sroF^Goe11^ into Δ6s.

Δ*aimP^6AA^* and chimeric aimP phage derivatives were cloned using the CRISPR-Cas9 vector pJoe899919. ^28^ Firstly, CRISPY-Web^29^ was used to identify appropriate candidate sgRNAs overlapping with the last six amino acids of AimP. These candidates were then synthesized as oligonucleotides with BsaI restriction sites and ligated onto the BsaI sites of pJoe8999. Subsequently, two 700bp fragments flanking aimP were amplified by PCR. AimP mutations were introduced as primer overhangs during the amplification of these fragments. We then used the NEBuilder HiFi DNA Assembly kit (New England Biolabs) to assemble the 700bp fragments with a SfiI-linearized, sgRNA-containing pJoe8999 plasmid. These constructed plasmids were transformed into *E. coli* on kanamycin (30 µg/ml) plates. Plasmids were screened with PCR and Sanger sequencing. Successful plasmid constructs were transformed into *B. subtilis* lysogens and plated on LB, supplemented with kanamycin (5 µg/ml) and 0.2% mannose, to induce Cas9 activity at 30°C for 24-48 hours. Colonies were then sub-streaked without antibiotics at 50°C to cure extra copies of the plasmid, sub-streaked to obtain single colonies at 42°C and screened for plasmid loss by testing for kanamycin sensitivity at 37°C. Successful clones were screened by colony PCR and Sanger sequencing. Lysogens containing successful mutants were then induced and lysogenized in new hosts to avoid bacterial host mutations. Briefly, 10uL of phage lysate was dispensed onto a freshly poured top agar lawn of *B. subtilis* Δ6s. The plates were grown overnight at 37°C, during which lysogenic colonies formed in the area where the lysate was spotted. The colonies were then swabbed and streaked onto a fresh LB plate to obtain single colonies. Single colonies were then restreaked to ensure a pure culture was obtained. Lysogens were screened by PCRs and Sanger sequencing, then sent for whole-genome sequencing to ensure no other mutations occurred in the phage during cloning.

To clone the tetracycline marker in WT Goe11 phages, PCRs were used to amplify the tetracycline cassette *tetR* with flanking sequences overlapping *yokI*. The PCR product was then transformed into Goe11 WT lysogen to replace *yokI*. Successful clones were screened using tetracycline plates, and verified with PCRs.

To generate reporter strains, the pJAL plasmid was developed by combination of the plasmids pAX01 and pDG1663 (Table S5). First, a synthetic DNA duplex containing N-terminal His tag and C-terminal streptavidin tag (Table S5) was inserted into plasmid pAX01 using Nebuilder Hifi DNA Assembly (BioEngland) commercial kit. The modified pAX01 integration cassette, containing the *ermR* gene, the xylose-inducible promoter Pxyl and both protein tags was amplified and used to substitute the integration cassette of pDG1663, *β-gal* and *spc* genes were removed. Both fragments were amplified by PCR with primers pAX_Cassette_pJAL_fwd and pAX_Cassette_pJAL_rev (Table S5) and ligated using Nebuilder Hifi DNA Assembly kit. Reporter strains carried two different insertion plasmids: pJAL, lacking tags, carrying the constitutive expression of *aimR* and pAND_101 carrying the *sfGFP* reporter cassette under control of the promoter regulated by AimR. All plasmids were cloned by combination of PCR fragments through Nebuilder Hifi DNA Assembly and constructs were verified by Sanger sequencing.

To produce recombinant AimR^13952^ genomic DNA from *B. subtilis* strain ATCC13952 was used for amplification of *aimR* gene from 13952 phage using primers ATCCPlic_FW and ATCCPlic_RV (Table S5). The PCR product was purified and cloned into pLicSGC1 plasmid using Ligation-Independent Cloning (LIC) system. The resulting plasmid, pLicSGC1-aimR^13952^ (Table S5) express full length AimR^13952^ merged with an N-terminal 6xHistag followed by a TEV protease cleaving site.

### Making conditioned media

The protocol to make conditioned media was adapted from Erez *et al.*^5^ Briefly, overnight cultures of *B. subitilis* Δ6s were diluted to OD600 0.05 in fresh minimal media and shaken at 180 rpm, 37°C until OD600 0.2. Then, 15 mL of culture was infected with phages at an MOI: 1. An uninfected control media was also prepared, to which no phage was added. The culture was further shaken at 180 rpm 37°C with durations varying by phage added (1.5 hours for Goe11, 13952 and Phi3T, 2.5 hours for Spbeta, 2.5 hours for control media). The varying durations were to ensure the media was filtered at the moment of maximum lysis by the phages, based on infection growth curves. The culture was then spun down at 4000 rpm for 10 min at 4 °C, and filtered with a 0.22 μm syringe filter, followed by filtration with a 3 kDa Amicon Ultra centrifugal filter (Milipore). Titration assays ensured no phages were in the media. The conditioned media were kept at 4 °C until use.

### Induction assays

Induction assays for synthetic AimP induction screens were conducted in LB media, while induction assays which compared WT, Δ*aimP^6AA^*, and chimeric phages were conducted in minimal media. This is because prior studies have reported that the differences in titres from adding synthetic peptide to WT phages are more pronounced in LB than minimal media, while differences in induction between WT and Δ*aimP^6AA^* phages are more pronounced in minimal media.^31^

To test inductions with synthetic peptides and the chimeric AimP phages, overnight cultures of lysogens were diluted to OD600 0.05 in the appropriate media. The media was supplemented with 0.1 mM MnCl2 and 5 mM MgCl2 and shaken at 180 rpm, 37°C until an OD600 value of 0.18. Synthetic AimP peptides were added to a concentration of 5 µM when indicated. Cultures were then shaken until OD600 0.2, after which mitomycin C (MC) was added to a final concentration of 0.5 μg/mL. Cultures in LB media were then shaken at 30°C, 80 rpm for three hours, and then left at room temperature without shaking for eight hours. Meanwhile, cultures in minimal media were shaken at 30°C, 80 rpm for 12 hours. Cultures were then 0.22 μm filtered and stored at 4°C for titration. To test mixed population inductions, the initial subcultures were grown in minimal media to OD600 0.2, then the lysogen cultures were then mixed in a ratio of 1:1, followed by the addition of MC to a final concentration of 0.5 μg/mL. The mixed cultures were then shaken at 30°C, 80 rpm for 12 hours, 0.22 μm filtered and stored at 4°C for titration.

A modified protocol was used to test inductions in conditioned media: after subculturing ΔaimP6AA lysogens in minimal media until OD600 0.2, 1 mL of culture was mixed with 1 mL of conditioned media. The mixture was then shaken at 180rpm, 37°C for 15 min, followed by the addition of MC to a final concentration of 0.5 μg/mL. Cultures were then shaken at 30°C, 80 rpm for two hours, followed by 0.22 μm filtration and storage at 4°C for titration. Shorter induction times were used compared to other induction experiments due to quick depletion of nutrients and AimP peptides in the spent conditioned media.

### Bacteriophage titration assays

*B. subtilis* Δ6s was the recipient for quantifying phages in the synthetic peptide screens, chimeric phages, and conditioned media inductions. Overnight cultures of *B. subitilis* Δ6s were diluted to OD600 0.05 in fresh LB media and shaken at 180 rpm, 37°C until an OD600 value of 0.2. 100 μL of bacterial culture were then mixed with 100 μL of lysate at the appropriate dilution for 10 min at room temperature. The mixture was then plated with 3 mL of 0.5 % top agar onto LB plates. The top agar and LB plates were both supplemented with 0.1 mM MnCl2 and 5 mM MgCl2. The top agar was left to dry for 20 min, followed by incubation at 37°C degrees overnight. Plaques were counted the next day.

To separately quantify Goe11 and 13952 in mixed population experiments, different titration recipients were used. First, cells expressing the phage master repressors *sroF^Goe11^* or *sroF^13952^* were tested to see if they can block infection by Goe11 or 13952, using media supplemented with 1 mM IPTG. Interestingly, while SroF^13952^ expression blocked only its cognate phage, SroF^Goe11^ expression blocked both phages, making it impossible to separately quantify the number of plaques formed by 13952. Therefore, we instead tested the ability of the phages to infect a mutant recipient strain Δ*ggaB* Δ6, lacking the genes required for the synthesis of a cell wall teichoic acid poly(glucosyl N-acetylgalactosamine 1-phosphate) [poly(GlcGalNAc 1-P)], which is produced by the ggaAB operon,^32^ and was reported to be necessary for infection of some *B. subtilis* phages.^33^ Infection by Goe11, as with phages Spbeta and Phi3T, required the presence of the ggaA, but was not necessary for the infection by phage 13952. Thus, a Δ*ggaB* Δ6 mutant was cloned and used as the titration recipient to quantify the number of plaques formed by 13952 and B. subtilis Δ6s expressing *sroF*^13952^ was used to quantify the number of Goe11 infective particles.

### Infection growth curves

To test the behavior of the chimeric phages in infection, overnight cultures of *B. subitilis* Δ6s were diluted to OD600 0.05 in fresh minimal media and shaken at 180rpm, 37°C until OD600 0.2. The culture was then diluted to OD600 0.05 and further grown to about OD600 0.18 to ensure exponential growth. Phages were then added to the culture at an MOI of 0.1. OD600 measurements were then taken on a 96-well plate reader for 6 hours.

### Lysogeny assays

For the infection ecological experiment, overnight cultures of 13952 lysogens were grown to OD600 of 0.2, and then infected with WT Goe11 encoding a tetracycline marker at MOI 1. At 1.5 hours, when maximum phage lysis occurs, the infection mixtures were plated on LB or LB with tetracycline plates to quantify the number of total cells and Goe11 lysogens respectively. 4 colonies were then picked from the tetracycline plate for whole genome sequencing, to verify that the colonies were indeed double lysogens.

### Reporter assays

Three overnight cultures of the reporter strain in parallel were diluted to OD600 0.05 in fresh LB medium and grown at 37°C and 230 rpm until OD600 reached 0.2. Then dilution was repeated in minimal media and when OD600 reached 0.2 again 100 µl of the cultures was transferred to a 96-well plate and combined with 100 µl of fresh minimal media supplemented with the analysed peptides at final concentration was 0.1 µM. In the case of Phi3T reporters IPTG at a final concentration of 1 µM was added since the construct lacks a constitutive promoter and an IPTG inducible promotor was introduced in the pAND_PPhi3T construct. The plate was incubated in Varioskan LUX (Thermo Scientific) microplate reader for 4 hours at 37°C with background shaking. Fluorescence was measured every 15 min, using 435/535 nm excitation/emission filters.

### Data visualization and statistical analyses

Differences in phages titres between groups were calculated with one-way ANOVA, followed by either Dunnett’s test or Tukey’s test using the stats package in R. Phage titres were log10 transformed. Experiments were visualized on R using the packages ggplot, and ggpubr.

### Constructing AimR-AimP clade2 database

The clade 2 AimR-AimP database was compiled by similarity searches with BLAST^34^ using the full length AimR^Spb^ against the NCBI reference proteins (refseq protein) database^35^ adjusted with default web server parameters an e-value threshold of 1×10^−40^. The more distant homologues were used as search models in successive BLAST rounds. To remove sequence redundancy the final set of sequences was filtered with Expassy Decrease Redundancy program (https://web.expasy.org/decrease_redundancy/) filtering to 99% of maximum similarity. In order to ensure that selected sequences belong to AimR clade2, a multiple sequence alignment (MSA) of the protein sequences of the AimRs was performed using ClustalW in Ugene (Unipro). The resulting MSA was used to create a maximum likehood phylogenetic tree using Treeviewer (https://treeviewer.org/) with 1000 bootstrap replicates. The selected clade2 contains 286 AimR sequences for which the immediately downstream gene corresponds to *aimP*. In the cases were the AimP ORF was not annotated, a fragment of 600 bases including the terminal nucleotides of each *aimR* gene was translated using the Expassy Translate Tool (http://web.expasy.org/translate/) and inspected to search AimP signature. The last six aminoacids of each AimP were considered as the mature peptide.

### Recombinant protein expression and purification

For AimR^13952^ production, a single colony of *E. coli* strain BL2_Codon plus (DE3) RIL (Agilent) carrying the pLicSGC1-aimR^13952^ plasmid was grown overnight at 37°C in 200ml of LB medium supplemented with 100 µg/ml of ampicillin and 33 µg/ml of chloramphenicol. The overnight culture was used at a 1:100 dilution to inoculate 6 one liter flasks of LB medium supplemented with ampicillin and chloramphenicol that were grown until an OD600 0.4, then the temperature was lowered to 20°C and IPTG to final concentration of 0.2 mM was added. After 16 hours of additional growth at 20°C, cells were harvested by centrifugation at 4.000 *g* for 20 min and the pellet was suspended in lysis buffer (25 mM Tris-HCl pH 8, 250 mM NaCl) and lysed by sonication on ice. Cell debris was removed by centrifugation at 10.000 *g* for 1 hour. The supernatant was loaded onto a 5 mL HisTrapFF (Cytiva), washed with lysis buffer and eluted with lysis buffer supplemented with 500 mM imidazole. Fractions containing the purest protein were pooled and digested with recombinant TEV protease fused to a Histag (50:1 molar ratio protein:TEV) and dialyzed against dialysis buffer (25mM Tris-HCl pH8, 250mM NaCl, 1mM β-mercaptoethanol and 0.5 mM DTT). In order to remove non digested protein and TEV, dialyzed sample was loaded in a 1 mL HisTrapFF (Cytiva). The flow though was collected, concentrated in a centrifugal filter unit (Amicon^TM^) and loaded in a Hi-Load Superdex 200 16/60 (GE Healthcare) gel filtration column equilibrated in lysis buffer. The purest fractions judged by SDS-PAGE were pooled, concentrated at 100 mg/ml and stored at −80°C. Typical yields were 25 mg recombinant protein/L of culture medium. AimR^Goe11^ and AimR^Phi3T^ were produced and purified as previously described.^16,17^

### Protein Crystallization and data collection

Crystallization assays were carried out at 21°C by vapour diffusion in hanging drops. Initial crystallization trials using commercial screens JBS I, JBS II (JENA Biosciences) and MIDAS (Molecular Dimensions) in 96-well plates (Swissci MRC2) were set up in the Cristalogenesis service of the IBV-CSIC, using a 10 mg/ml dilution of the protein in lysis buffer and mixing equal volumes of protein and mother solutions. The apo form of AimR^13952^ crystallized in 0.1 M Lithium sulfate, 0.1M HEPES 7.0 and 30 % w/v Polyvinylpyrrolidone. To obtain the AimR-AimP complexes, peptide was added to the protein solution at a final concentration of 1 mM and incubated for 10 min before setting up the crystallization plates. The AimR-AimP^13952^ crystallized in 30% PEG 3000, 0.1M CHES pH 9.5. The AimR^13952^-AimP^Goe11^ crystallized in 0.2 M Lithium sulfate, 15% PEG 8000. AimR^Goe11^-AimP^13952^ crystallized in 15% pentaeritritol ethoxilate, 3% jeffamine, 0.2M Potassium Acetate, 0.1 M MES. Crystals grew in 2-7 days and were directly flash frozen in liquid nitrogen. Diffraction data from single crystals was collected at Xaloc beamline (Alba Synchrotron) at 100°K. Data sets were processed with XDS^36,37^ and reduced using Scala (CCP4). ^37^ The data-collection statistics for data sets used in structure determination are shown in Table S2.

### Phase determination, model building and refinement

All structures were determined by molecular replacement with Phaser^38^ using AimR^Goe11^ coordinates (7Q0N^17^) as search model. Interactive cycles of manual model building using COOT^39^ and computational refinement with Refmac^40^ were used to generate the final model. Programs used are included in CCP4 Cloud package.^41^ Data collection and refinement statistics are summarized in Table S2.

### Thermal Shift Assay

The thermal shift assay was conducted in a 7500 Fast Real time PCR System (Applied Biosystems) as previously described.^42^ Briefly, samples of 20 μL buffer (20 mM Tris pH 8 and 250 mM NaCl) with 5XSypro Orange (Sigma-Aldrich) and 20 μM of protein were loaded in 96-well PCR plates. Peptides to 0.5 μM final concentration were added to the mixture when stabilization in peptide presence was evaluated. Samples were heated from 25 to 85°C in steps of one degree. Melting temperatures (Tm) were calculated plotting the fluorescent intensity versus temperature and integrating with GraphPad Prism software using a Boltzmann model.

### EMSA assays

Native polyacrylamide gel electrophoresis was used to check AimR binding to its operator were done as previously described.^17^ Double strand DNA primer probes were purchased from IDT. Briefly, AimR at 0.5 µM and DNA at 10 ng/ µl were mixed in EMSA buffer (50 mM Tris pH8, 250 mM NaCl). Samples were incubated at room temperature for 10 min before loading. For peptide inhibition evaluation, protein was preincubated with 0.5 µM peptide for 10 min before DNA addition. Electrophoresis was performed in 8% polyacrylamide gels in Tris-Borate-EDTA (TBE) buffer for about 110 min at 100 V at 4 °C.

### Isotermal Titration Calorimetry (ITC)

A Nano ITC Low Volume (TA instruments) was used to perform ITC and calculate the dissociation constant of AimR^Goe11^, AimR^13952^ and AimR^Phi3T^ against GIVRGA, GVVRGA and SAIRGA peptides. Protein at 15 µM was assayed against peptides at 150-200 µM. In all cases, both proteins and peptides were diluted in lysis buffer. The experiment was performed at 25°C. The NanoAnalyze software (TA Instruments) was used to integrate, correct and analyze the data using a single-site binding model.

### Size Exclusion Chromatography with Multi-Angle Light Scattering (SEC-MALS)

SEC-MALS experiments were performed using a Wyatt DAWN HELEOS-II MALS instrument and a Wyatt Optilab rEX differential refractometer (Wyatt) coupled to an AKTA pure system (GE Heralthcare). 20 µL of protein at 5 mg/mL were injected on a KW-803 (Shodex) column equilibrated in lysis buffer. When testing AimR-AimP^13952^ oligomeric state, the protein sample was incubated with 1mM peptide for 10 min before injection and the running buffer was supplemented with peptide at a final concentration of 1 μM. The Astra 7.1.2 software from the manufacturer was used for acquisition and analysis of the data.

## Acknowledgments

We thank the IBV-CSIC Crystallogenesis Facility for protein crystallization screenings. Data collection experiments for structural data were carried out at XALOC beamline at ALBA synchrotron (Cerdanyola del Valles, Spain). The structural results reported in this article derive from measurements made at the synchrotron DLS ALBA (Cerdanyola del Valles, Spain) and ESRF (Grenoble, France). Data collection experiments for the structures reported in the manuscript were carried at XALOC beamline at ALBA synchrotron. X-ray diffraction data collection was supported by block allocation group (BAG) ALBA Proposal 2023077633. We acknowledge the ALBA synchrotron for provision of beam time and we would like to thank beamline staff for assistance. Some figures in this manuscript have been created with Biorender.com.

This work was supported by grants PID2022-137201NB-I00 from Spanish Government (Ministerio de Ciencia e Innovación), CIPROM/2023/30 from Valencian Government and the European Commission NextGenerationEU fund (EU 2020/2094), through CSIC’s Global Health Platform (PTI Salud Global) to A.M; A.M. and J.R.P are funded by European Research Council Grant 101118890 (TalkingPhages).

## Data Availability

All data are available from the authors upon request. Coordinates and structure factors for AimR^13952^, AimR-AimP^13952^, AimR^13952^-AimP^Goe11^ and AimR^Goe11^-AimP^13952^ have been deposited in the Protein Data Bank with access codes 9F36, 9F82, 9F9R and 9FKU, respectively. All relevant accession codes and identifiers are provided within the manuscript. Figures do not have associated raw data. No restrictions on data availability.

