## Supplemental Table S5 for "Phages communicate across species to shape microbial ecosystems": Gallego_et. al-Supplemental-information.pdf

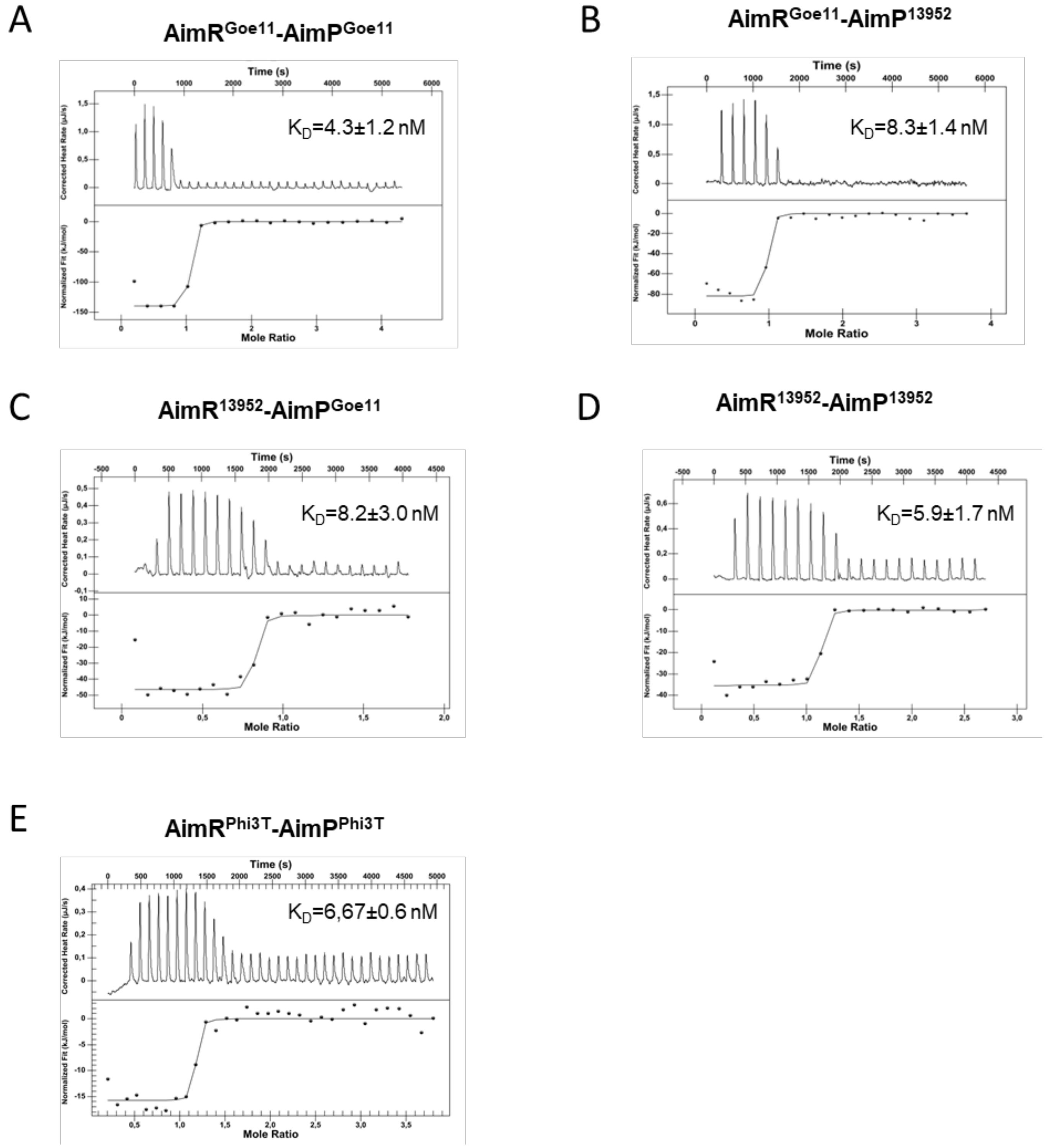


**Figure S1: ITC Graphics. In vitro data shows that $AimR^{Goe11}$ and $AimR^{13952}$ can bind non canonical peptides.** Binding affinities measured by ITC measurement for $AimR^{Goe11}$-$AimP^{Goe11}$ (A), $AimR^{Goe11}$-$AimP^{13952}$ (B), $AimR^{13952}$-$AimP^{Goe11}$ (C) and $AimR^{13952}$-$AimP^{13952}$ (D) and $AimR^{Phi3T}$-$AimP^{Phi3T}$. Thermograms are adjusted to one-binding site model and the $K_{Ds}$ are shown in a box in each graphic.

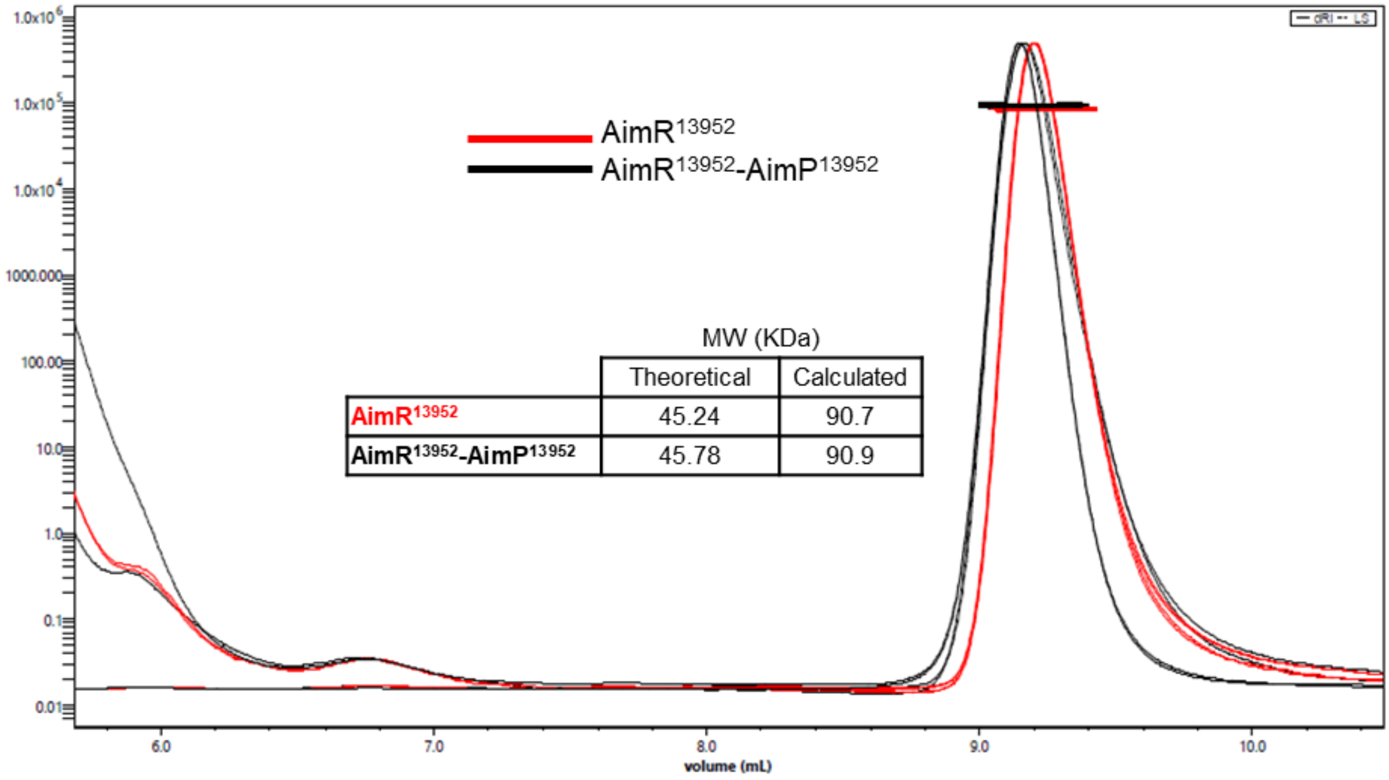


| | Theoretical | Calculated |
|---|---|---|
| AimR13952 | 45.24 | 90.7 |
| AimR13952-AimP13952 | 45.78 | 90.9 |

**Figure S2. AimR$^{13952}$ oligomeric state.** Size exclusion chromatography multi-angle light scattering (SEC-MALS) chromatograms of AimR$^{13952}$ in absence (red) and presence (black) of AimP$^{13952}$ peptide (GVVRGA). Chromatograms show the readings from the light scattering and refractive index detectors. The vertical axis represents the molecular mass. The horizontal curves represent the calculated molecular masses. In both cases the molecular weight calculated corresponds to a dimer (~90kDa).

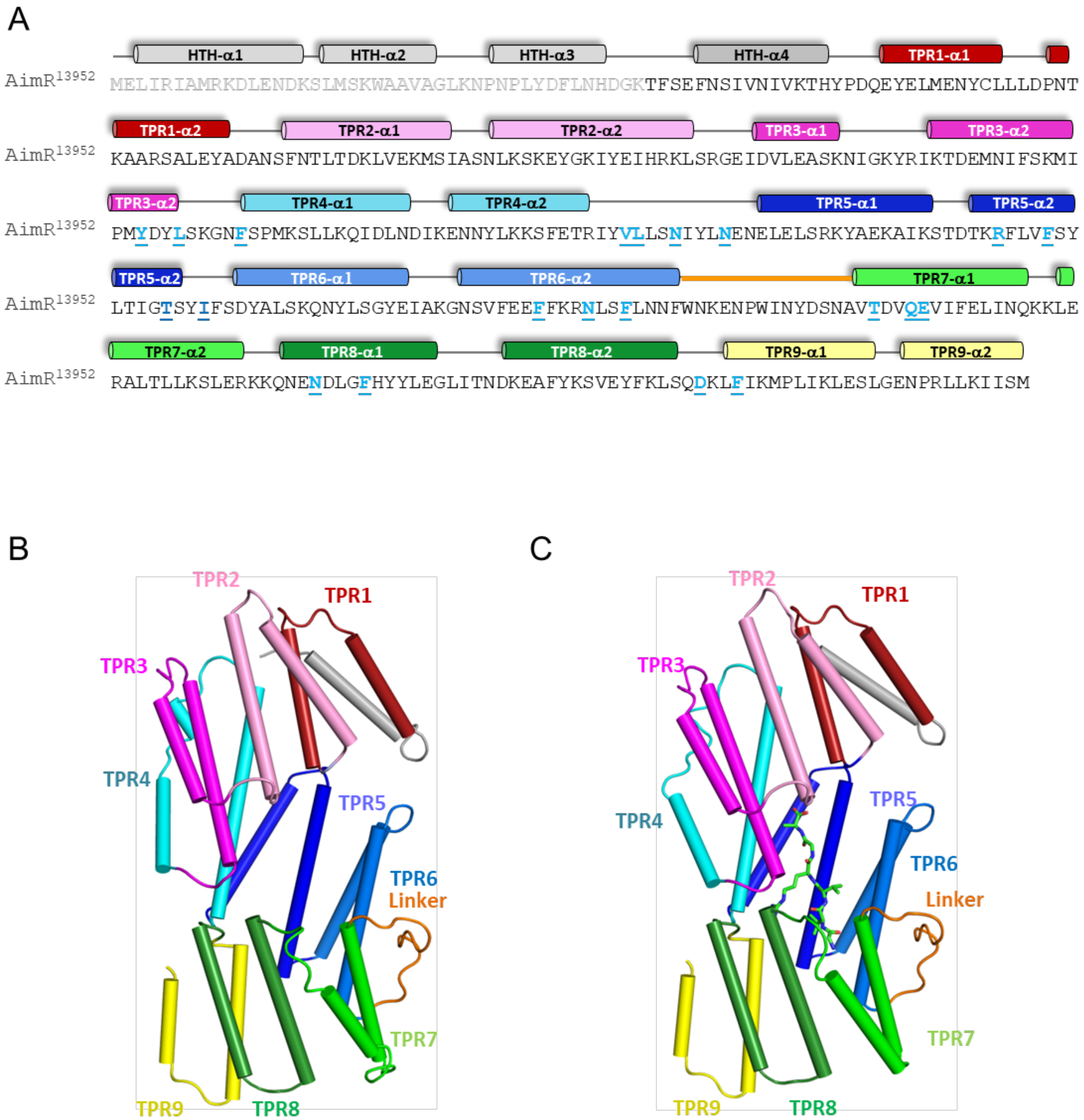


**Figure S3: Sequence and TPR domain organization in AimR$^{13952}$.** (A) AimR$^{13952}$ sequence with residues involved in peptide binding underlined, highlighted in bold and coloured in blue. Structural elements are shown above and labelled by helices for HTH and TPR domains. (B) Cartoon representation of AimR$^{13952}$ crystal structure, with a-helices coloured in the same colour code used used in A. (C) Cartoon representation of AimR$^{13952-}$ AimP$^{13952}$ complex, with AimP shown as green sticks.

A

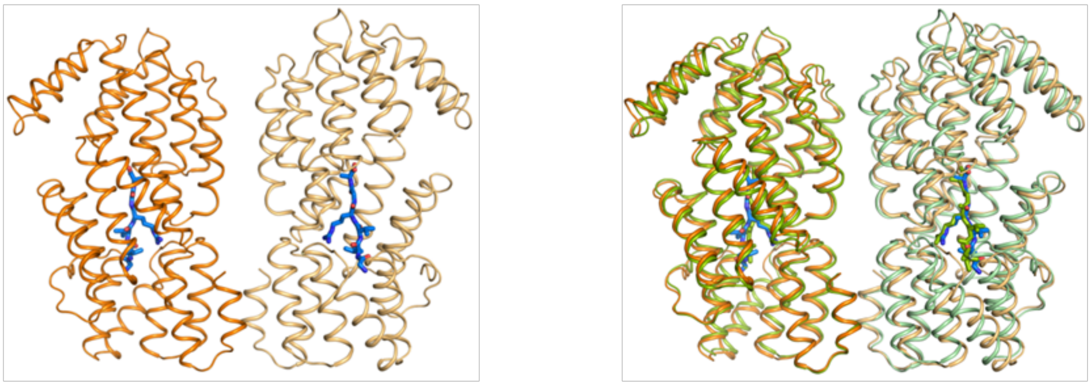


B

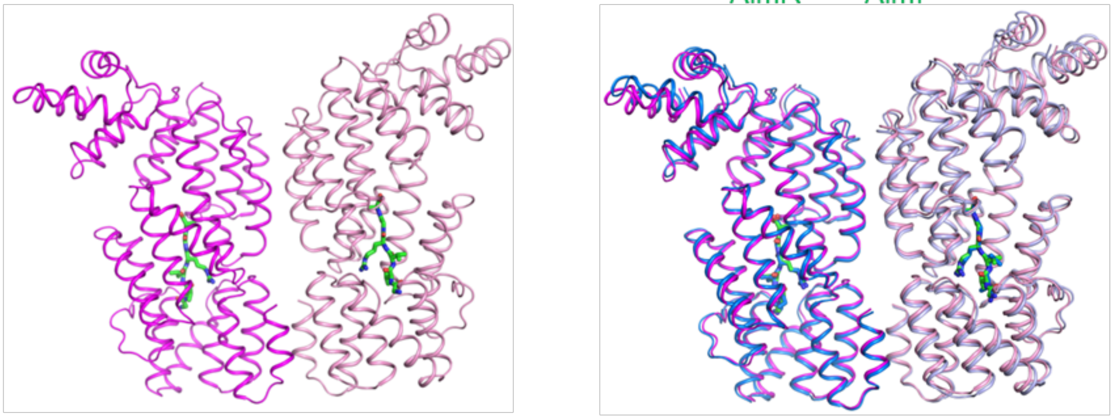


C

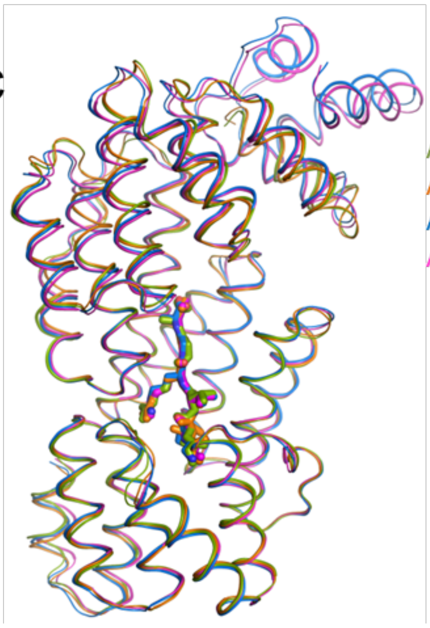


| | | **AimR$^{13952}$** | | **AimR$^{Goe11}$** | |
|---|---|---|---|---|---|
| | | **GVVRGA** | **GIVRGA** | **GVVRGA** | **GIVRGA** |
| **AimR$^{13952}$** | **GVVRGA** | 0 | 0.51 | 0.74 | **0.76** |
| | **GIVRGA** | | 0 | **0.86** | **0.84** |
| **AimR$^{Goe11}$** | **GVVRGA** | | | 0 | 0.46 |
| | **GIVRGA** | | | | **0** |

**Figure S4: Superposition of AimR-AimP complexes.** A) Cartoon representation of AimR$^{13952}$-AimP$^{Goe11}$ (left) and superposition of AimR$^{13952}$ in complexes with its cognate and non cognate AimPs. B) Cartoon representation of AimR$^{Goe11}$-AimP$^{13952}$ (left) and superposition of AimR$^{Goe11}$ complexes with its cognate and non cognate AimPs (right). In both figures AimPs are shown in sticks, and colored in blue for AimP$^{Goe11}$ or green for AimP$^{13952}$ C) Superposition of the four AimR-AimP combinations (left) and table showing Root Mean Square Deviations (RMSD) for monomer superposition in Å (right).

**Table S2.** Number and percentage of proposed mature peptides found in clade 2.

| Proposed mature peptide | Total number | % |
|---|---|---|
| GFGRGA | 57 | 19.9 |
| SASRGA | 42 | 14.7 |
| GMPRGA | 32 | 11.2 |
| GFTVGA | 25 | 8.7 |
| SAIRGA | 19 | 6.6 |
| SIIRGA | 17 | 5.9 |
| NPGRGA | 12 | 4.2 |
| DPGRGG | 9 | 3.1 |
| GFGHGA | 9 | 3.1 |
| GVVRGA | 9 | 3.1 |
| SIGHGA | 9 | 3.1 |
| SPSRGA | 8 | 2.8 |
| GFPRGA | 5 | 1.7 |
| GIVRGA | 5 | 1.7 |
| TIGRGG | 5 | 1.7 |
| AIGNGG | 4 | 1.4 |
| AMGNGG | 4 | 1.4 |
| ATIGRG | 3 | 1.0 |
| GDGGRP | 2 | 0.7 |
| GMGRGA | 2 | 0.7 |
| GMVRGA | 1 | 0.3 |
| GTRPPS | 1 | 0.3 |
| KPGIGG | 1 | 0.3 |
| NPAYME | 1 | 0.3 |
| NPGRHA | 1 | 0.3 |
| RPGVGA | 1 | 0.3 |
| SMVRGA | 1 | 0.3 |
| ∅ | 1 | 0.3 |
| **Total** | 286 | 100 |

**Table S3.** Data collection and refinement statistics

| | AimR$^{13952}$ | AimR$^{13952}$- AimP$^{13952}$ | AimR$^{13952}$- AimP$^{Goe11}$ | AimR$^{Goe11}$- AimP$^{13952}$ |
|---|---|---|---|---|
| **Data collection** | | | | |
| Space group | $P2_12_12$ | $P2_1$ | $P_1$ | $C222_1$ |
| Cell dimensions | | | | |
| *a*, *b*, *c* (Å) | 82.99, 150.77, 67.33 | 67.34, 82.96, 146.86 | 44.63, 67.04, 144.62 | 69.77, 203.58, 143.07 |
| α, β, γ (°) | 90, 90, 90 | 90, 90, 90 | 90.20, 94.25, 107.48 | 90, 90, 90 |
| Resolution (Å) | 150.8-2.3 (2.3-2.36)* | 146.8 (3.1) (3.1-3.18) | 141.6 (2.5) (2.5-2.64) | 48.6 (1.9) (1.9-1.95) |
| $R_{pym}$ | 0.02 (0.49) | 0.06 (0.24) | 0.059 (0.41) | 0.029 (0.455) |
| $I$ / σ$I$ | 16.9 (1.4) | 8.7 (3.8) | 8.2 (1.8) | 16.2 (1.7) |
| Completeness (%) | 100 (99.9) | 99.8 (99.9) | 83.9 (49.5) | 100 (100) |
| Redundancy | 12.5 (6.7) | 6.7 (6.8) | 3.4 (2.9) | 6.7 (6.3) |
| **Refinement** | | | | |
| Resolution (Å) | 2.3 | 3.1 | 2.5 | 1.9 |
| No. reflections | 36403 (2650) | 29584 (4750) | 34420 (1721) | 80454 (7816) |
| $R_{work}$ / $R_{free}$ | 0.19(0.36)/ 0.25(0.46) | 0.23 (0.27)/0.28 (0.36) | 0.20 (0.36)/0.25(0.40) | 0.17 (0.32)/0.21 (0.33) |
| No. atoms | | | | |
| Protein | 5700 | 11464 | 11468 | 6443 |
| Ligand/ion | 5 | | | 36 |
| Water | 66 | 24 | 58 | 396 |
| *B*-factors | | | | |
| Protein | 73.8 | 80.6 | 38.3 | 40.8 |
| Ligand/ion | 107.5 | | | 63.5 |
| Water | 60.5 | 46.0 | 18.4 | 41.1 |
| R.m.s. deviations | | | | |
| Bond lengths (Å) | 0.01 | 0.014 | 0.012 | 0.013 |
| Bond angles (°) | 1.82 | 2.35 | 2.06 | 1.9 |
| **PDB Access Code** | 9F36 | 9F82 | 9F9R | 9FKU |

*Values in parentheses are for highest-resolution shell.

**Table S4: AimR-AimP interactions.** The AimR residues interacting with the AimP main- and side-chain in each complex are coloured in orange and black, respectively. Non homologous AimR residues are shadowed in yellow.

| Receptor | | AimR$^{13952}$ | | AimR$^{Goe11}$ | |
|---|---|---|---|---|---|
| **AimP Position** | **AimP Residue** | **GVVRGA** | **GIVRGA** | **GVVRGA** | **GIVRGA** |
| **P1** | **G** | T296 | T296 | T296 | |
| | | Q299 | Q299 | Q299 | Q299 |
| | | E300 | E300 | E300 | E300 |
| **P2** | **V/I** | | | | T239 |
| | | | | | F276 |
| | | F269 | F269 | | |
| | | F333 | F333 | F333 | F333 |
| | | F363 | F363 | (S363)* | (S363)* |
| **P3** | **V** | F269 | F269 | F269 | F269 |
| | | N273 | N273 | N273 | N273 |
| **P4** | **R** | F167 | F167 | F167 | F167 |
| | | N206 | N206 | N206 | N206 |
| | | N329 | N329 | N329 | N329 |
| | | D360 | D360 | D360 | D360 |
| **P5** | **G** | L162 | | | |
| | | N202 | N202 | N202 | N202 |
| | | F232 | F232 | F232 | F232 |
| **P6** | **A** | | Y159 | Y159 | Y159 |
| | | L162 | | | |
| | | V198 | V198 | V198 | V198 |
| | | R228 | R228 | R228 | R228 |

* S363 from AimR$^{Goe11}$ is shown to notice the residue change between AimRs but not interact with AimP.
